# Dominant negative effects on H3K27 methylation by Weaver syndrome-associated EZH2 variants

**DOI:** 10.1101/2023.06.01.543208

**Authors:** Orla Deevy, Craig Monger, Francesca Matrà, Ellen Tuck, Eric Conway, Mihaly Badonyi, Darragh Nimmo, Simona Rodighiero, Qi Zhang, Chen Davidovich, Joseph A. Marsh, Diego Pasini, Adrian P. Bracken

## Abstract

Heterozygous missense mutations in *EZH2* cause Weaver syndrome (WS), a developmental disorder characterized by intellectual disability and overgrowth. *EZH2* encodes the enzymatic subunit of Polycomb Repressive Complex 2 (PRC2), which mediates mono-, di-, and tri-methylation of histone H3 lysine residue 27 (H3K27me1/2/3). Although the functional characterization of most WS-associated EZH2 variants is lacking, they are presumed loss of function. However, the dearth of reported early truncating mutations in *EZH2* led us to hypothesise that a dominant negative mutational mechanism may underlie the development of WS. To test this, we performed a detailed structural analysis of all known WS-associated EZH2 variants, which provided initial support that they are dominant negative. Next, we isogenically modelled 10 representative WS-associated EZH2 variants in embryonic stem cells and showed they induce global reductions in H3K27me2 and H3K27me3 with concomitant global increases in H3K27me1, H3K27ac, and chromatin decompaction. Importantly, the reductions in H3K27me2/3 methylation revealed a pattern of dominant-negative interference to PRC2 activity. Comparative analysis of a gain-of-function EZH2 variant causing growth restriction highlighted the reciprocal nature of the chromatin changes in these opposing growth syndromes. Our findings detail the molecular effects of developmental-syndrome-associated EZH2 variants in cells and implicate imbalanced landscapes of H3K27 modification in their pathology.

## Introduction

Polycomb Repressive Complex 2 (PRC2) is a highly conserved chromatin regulatory complex essential for lineage specification and cellular memory in multicellular organisms (Margueron and Reinberg 2011; Schuettengruber et al. 2017; Laugesen et al. 2019). Mammalian PRC2 consists of three core subunits – SUZ12, EED, and either the EZH1 or EZH2 histone methyltransferase (HMTs) – which together catalyse the deposition of up to three methyl groups on histone H3 at lysine 27 (H3K27). This trimeric core of PRC2 interacts with accessory proteins that facilitate its recruitment to chromatin and modulate its methyltransferase activity (Healy et al. 2019; Laugesen et al. 2019; Glancy et al. 2021). Both core and accessory PRC2 components are essential for normal mammalian development (Deevy and Bracken 2019). PRC2 broadly deposits H3K27me3 across transcriptionally silent developmental gene promoters (Boyer et al. 2006; Bracken 2006; Lee et al. 2006). The H3K27me3 modification allosterically activates PRC2 via specific interactions with EED that promote further PRC2 binding to chromatin (Lee et al. 2018). H3K27me3 contributes to the recruitment of the canonical Polycomb Repressive Complex 1 (cPRC1) via specific interaction with its constituent CBX proteins (Bracken et al. 2019; Glancy et al. 2023). This cPRC1 is in turn thought to promote chromatin compaction and transcriptional repression (Bracken et al. 2019). The functions of PRC2-mediated H3K27me1 and H3K27me2 are less well understood. While H3K27me1 is localised along actively transcribed gene bodies (Ferrari et al. 2014; Lee et al. 2015), H3K27me2 is extensively deposited throughout intergenic regions of the genome (Conway et al. 2015; Streubel et al. 2018). The broad deposition profile of H3K27me2 prompted us to propose it could function as a repressive “blanket” to counter accumulation of H3K27 acetylation (H3K27ac), a histone modification that is enriched at transcriptionally active regions and mutually exclusive from H3K27me2/3 on a given histone tail (Ferrari et al. 2014; Conway et al. 2015; Lee et al. 2015).

Heterozygous mutations in the *EZH2, EED* and *SUZ12* genes cause a triad of congenital overgrowth and intellectual disability syndromes with highly overlapping clinical presentations – Weaver syndrome (MIM #277590; WS), Cohen-Gibson syndrome (MIM #617561; COGIS), and Imagawa-Matsumoto syndrome (MIM #618786; IMMAS), respectively. These PRC2-related overgrowth syndromes are rare, with collectively less than 100 cases reported to-date (Cyrus et al. 2019a; Cyrus et al. 2019b; Hetzelt et al. 2021). However, it is likely that their true incidence is underestimated as a result of their very wide clinical presentation, which most often includes pre- and post-natal overgrowth, tall stature, advanced bone age, variable intellectual disability, and a subtle but characteristic facial appearance (Tatton-Brown and Rahman 2013; Cyrus et al. 2019a). WS is the most extensively studied PRC2-related overgrowth syndrome, with *EZH2* first reported as the causative gene in 2011 and approximately 70 affected individuals reported to date (Tatton-Brown et al. 2011; Gibson et al. 2012). Most cases of WS are *de novo*, but the few familial cases exhibit an autosomal dominant inheritance pattern (Fryer et al. 1997; Proud et al. 1998; Tatton-Brown et al. 2013). Notably, almost all reported WS-associated *EZH2* mutations are missense variants, with the majority affecting its catalytic SET domain (Tatton-Brown et al. 2013). Rare truncating nonsense and frameshift mutations have also been reported though, intriguingly, they all occur within the last exon of *EZH2* and are therefore likely to escape nonsense-mediated decay at the mRNA level; in other words, an almost-full-length truncated EZH2 variant will be expressed (Hentze and Kulozik 1999; Imagawa et al. 2017). The lack of early truncating mutations in more N-terminal exons of *EZH2* suggests haploinsufficiency is unlikely to be the major mutational mechanism of WS. Regardless, WS-associated EZH2 variants are presumed loss-of-function hypomorphs due to the reduced HMT activity demonstrated for 10 WS-associated EZH2 variants studied to date (Cohen et al. 2016; Imagawa et al. 2017; Lee et al. 2018; Lui et al. 2018; Jani et al. 2019; Gao et al. 2024). Taking these pieces of genetic and functional evidence together, we speculate that WS-associated EZH2 variants are indeed partial loss of function but with potential dominant negative effects.

Thus far, functional molecular studies of these Weaver WS-associated EZH2 variants have employed *in vitro* HMT assays, DNA methylation profiling, or assays of global H3K27 methylation (Cohen et al. 2016; Lui et al. 2018; Jani et al. 2019; Choufani et al. 2020; Gao et al. 2024). However, these *in vitro* assays and global assessments of H3K27 methylation levels do not capture the complexity of the Polycomb system. Namely, they fail to delineate the three different H3K27 methylation states in cells — both genome wide and on Polycomb target genes — and they do not consider the possible downstream consequences on H3K27ac, cPRC1 recruitment, chromatin compaction, or target gene expression. Given that PRC2 function is frequently deregulated in cancer, with both activating and inactivating mutations in *EZH2*, a key motivation for a thorough molecular study of WS-associated EZH2 variants is the fact that this information may help clinicians more accurately predict what cancers their affected patients may be predisposed to (Conway et al. 2015; Laugesen et al. 2016). As such, we wished to test our hypothesis that WS-associated EZH2 variants may have dominant negative effects.

Here we detail the consequences of WS-associated EZH2 variants on the genome-wide functions of PRC2 in mouse embryonic stem cells (mESCs). We show that WS-associated EZH2 variants are hypomorphs that can dominant negatively impair PRC2-mediated H3K27 methylation, which could explain the lack of early truncating mutations in individuals with WS. We confirm that heterozygous WS-associated EZH2 variants reduce intergenic H3K27me2/3 with corresponding increases in H3K27ac and impaired chromatin compaction. Finally, a comparative analysis of WS-versus growth restriction-associated EZH2 variants highlighted the reciprocal chromatin changes potentially underpinning their opposing aberrant growth phenotypes. Taken together, our results shed new light on the molecular aetiology of WS and highlight how the precise equilibrium of H3K27 methylations is crucial for the regulation of growth during development.

## Results

### Weaver syndrome-associated EZH2 variants are predicted to be non-loss-of-function

We sought to perform a functional investigation of EZH2 variants associated with Weaver syndrome. We first performed an extensive literature review identifying 67 individuals diagnosed with Weaver syndrome who were reported to have a coding mutation in EZH2 (Fig. 1A, Table 1). Among this group, 61 individuals presented with missense variants affecting 34 different residues within EZH2, 3 individuals had in-frame indel mutations, and 3 individuals had C-terminal truncating mutations that are expected to produce a near-full-length protein product. Supporting impaired HMT activity as a central mechanism in WS, the single amino acid variants tend to cluster around the SET and SRM domains of EZH2, which together interface with the histone H3 tail substrate to mediate H3K27 methylation (Fig. 1B).

**Table 1.**
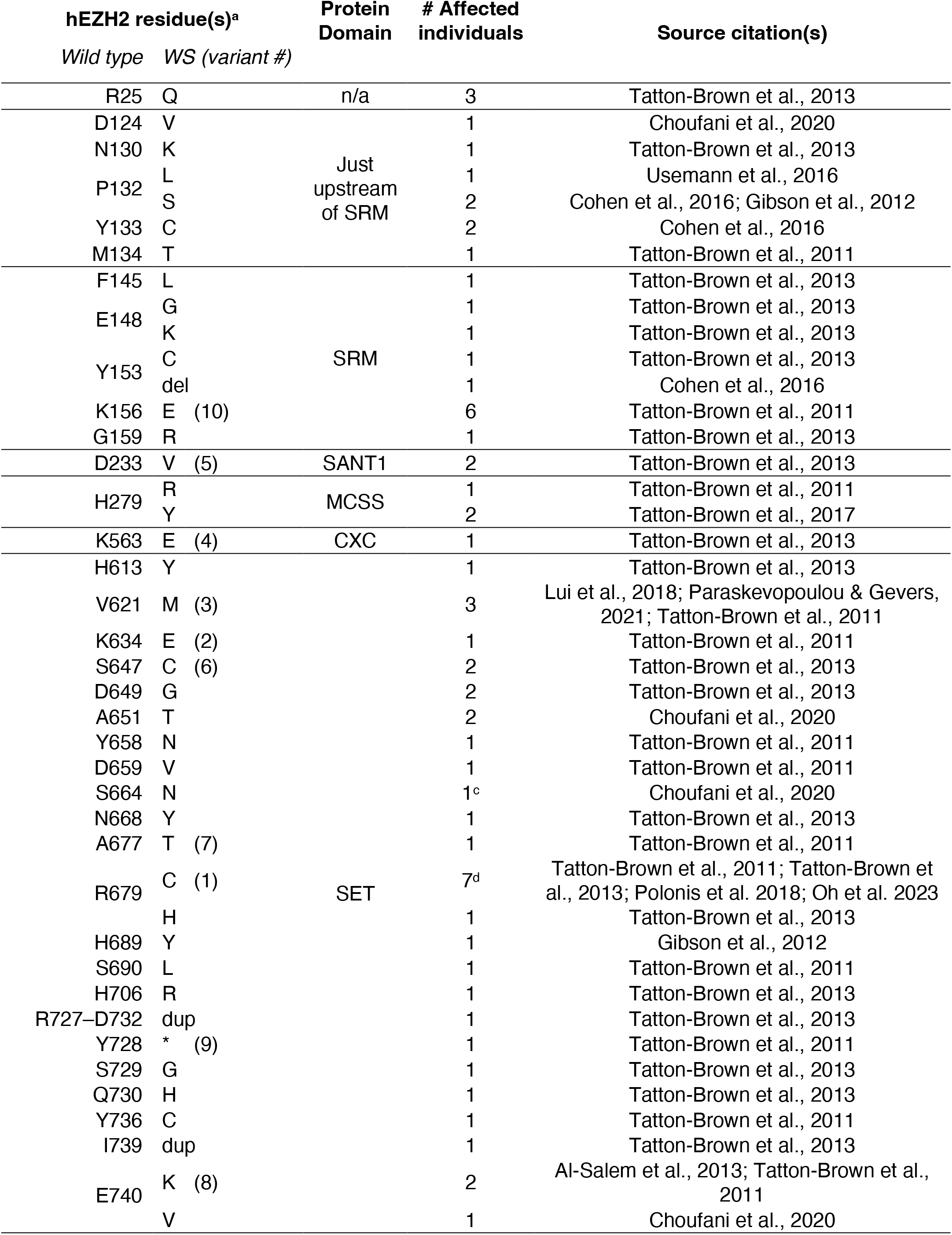
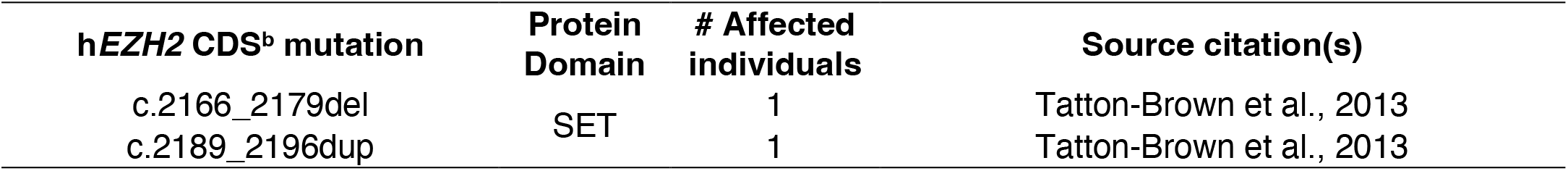
All Weaver syndrome-associated EZH2 variants reported to date. ^a^hEZH2 residue in isoform 1 (Q15910-1) ^b^CDS variants per HGVS nomenclature correspond to consensus coding sequence (CCDS56516.1) ^c^Choufani et al. report a father and child carrying the same pathogenic *EZH2* variant, where only the child is clinically affected; only the child is included in the number of affected individuals. ^d^Polonis et al. report a father and child carrying the same pathogenic *EZH2* variant, where only the father exhibits classic features of Weaver syndrome; only the father is included in the number of affected individuals. *Abbreviations: CDS, coding sequence; del, deletion; dup, duplication; h, human; n/a, not applicable; WS, Weaver syndrome; #, number*

**Figure 1.**
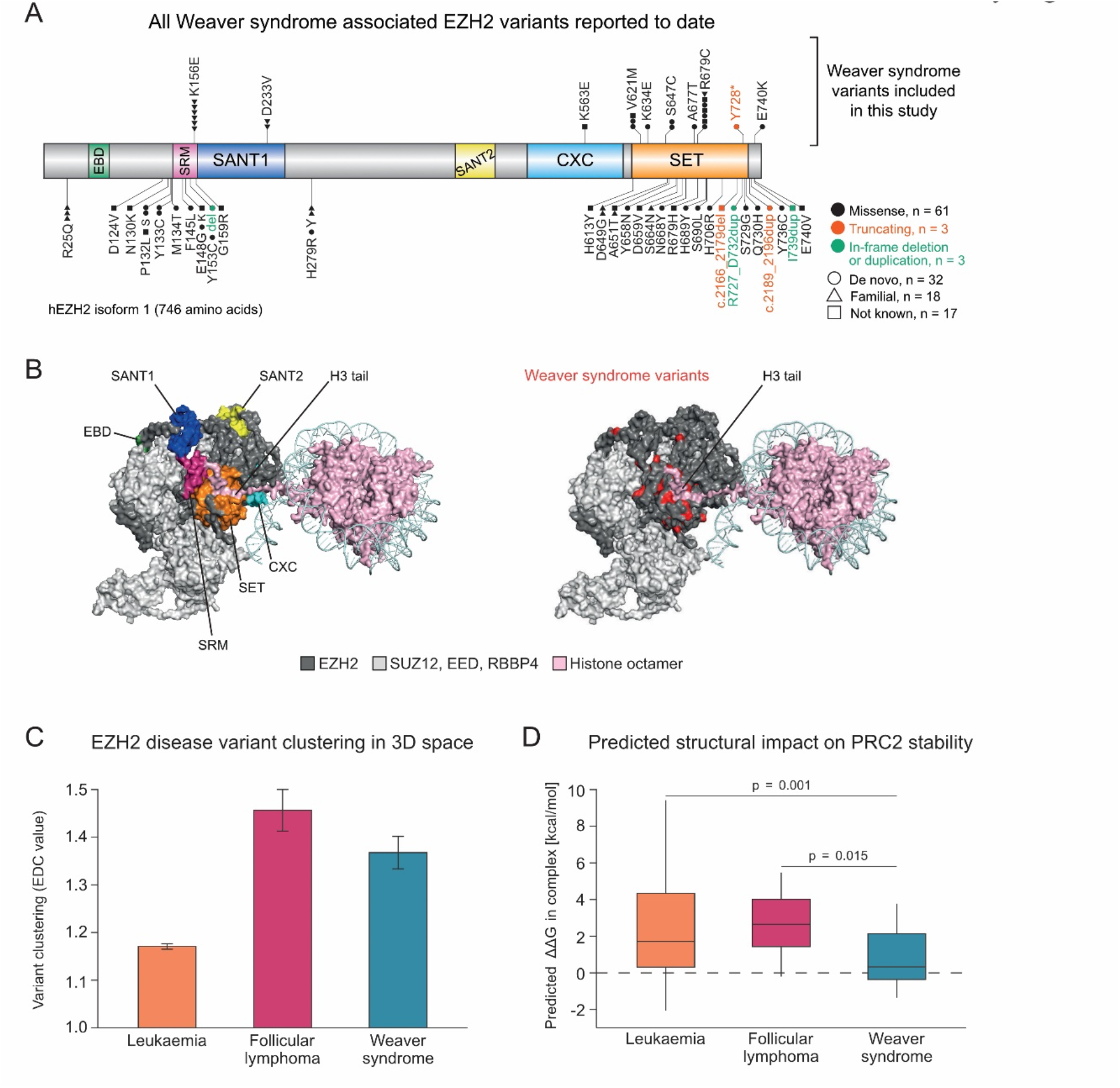
Weaver syndrome associated EZH2 variants cluster in domains proximal to the histone H3 tail substrate. **A**. Schematic of human EZH2 protein (isoform 1 (Q15910-1) summarising the distribution of coding variants reported in Weaver syndrome, highlighting those included in this study; Table 1 provides the source citations. Missense variants are indicated in black, nonsense variants in red, and in-frame duplications in green. Each shape represents an affected individual with a pathogenic mutation in EZH2 that is either *de novo* (circle), familial (triangle), or of unknown inheritance status (square). **B**. Cryo-EM structure of PRC2.2 bound to a H2AK119ub1-modified nucleosome, with EZH2 shown in dark grey, and EED and SUZ12 in light grey for clarity (PDB: 6WKR, Kasinath et al., 2021). Left: EZH2 domains are highlighted as per panel (a) above: EBD (light green), SRM (hot pink), SANT1 (dark blue), SANT2 (yellow), CXC (cyan) and SET (orange). Right: Amino acids affected by Weaver syndrome mutations are highlighted in red. Nucleosome DNA is shown in pale cyan and the histone octamer is shown in light pink. **C**. Graphical representation of the 3D clustering of disease variants associated with leukaemia, follicular lymphoma, and Weaver syndrome. The bars represent the raw EDC values of missense variants associated with the different phenotype groups. Higher values indicate stronger clustering within the 3D structure of EZH2-PRC2. Bar, raw EDC; error bars, standard deviation obtained with jackknife resampling. **D**. Predicted structural impacts (ΔΔG) on the EZH2:PRC2 complex structure of missense variants associated with different disease phenotypes. ΔΔG of missense mutations associated with the different phenotype groups. P-values were calculated with a two-sided Wilcoxon rank-sum test. Box, 25th and 75th percentiles; middle line, median; whiskers, 1.5 times the interquartile range. *Abbreviations: EBD, EED-binding domain; EDC, Extend of Disease Clustering; SRM, stimulatory response motif; 3D, three-dimensional; ΔΔG, the change in Gibbs free energy of folding*.

Interestingly, Gerasimavicius et al. (2022) showed that dominant-negative mutations are highly enriched at protein interfaces. They also demonstrated that the degree to which missense variants cluster in three-dimensional (3D) protein structures and their effects on complex stability correspond well to the underlying molecular disease mechanism. Loss-of-function variants tend to show weak structural clustering and to disrupt protein structure, whereas non-loss-of-function (i.e., gain-of-function and dominant-negative) variants tend to be highly clustered with mild effects on protein structure (Gerasimavicius et al. 2022). We compared these features among WS-associated missense EZH2 variants versus known loss-of-function and gain-of-function oncogenic EZH2 variants from leukaemia and follicular lymphoma, respectively (Table 2). Using the crystal structure of EZH2 in the PRC2 complex (Justin et al. 2016), we found that WS and follicular lymphoma variants strongly clustered relative to leukaemia variants (Fig. 1C). Next, we estimated the change in the free energy of folding (ΔΔG) associated with each pathogenic missense variant, where higher ΔΔG values indicate variants that are more likely to disrupt the structural stability of PRC2. Interestingly, while we found that leukaemia and lymphoma variants tended to be highly disruptive to protein structure, WS variants were predicted to be much less structurally damaging (Fig. 1D). To summarise, (1) leukaemia variants showed low clustering and were highly structurally damaging, consistent with a loss-of-function mechanism, (2) lymphoma variants were highly clustered and also highly disruptive to protein structure, consistent with gain-of-function mutations that act via localised structure damage (Backwell and Marsh 2022), and (3) the WS variants were highly clustered and structurally mild, strongly supportive of an underlying non-loss-of-function mechanism. Although this protein structural analysis alone is not able to distinguish between dominant-negative and gain-of-function effects, it is striking that WS variants are clearly distinct at the protein structural level from both the loss-of-function leukaemia variants and gain-of-function lymphoma variants.

**Table 2.** EZH2 variants in Weaver syndrome, lymphoma and leukaemia used in variant effect prediction analysis in Figure 1. *Abbreviations: WS, Weaver Syndrome*.

|  | WS | Lymphoma | Leukaemia |  |  |  |  |
| --- | --- | --- | --- | --- | --- | --- | --- |
| EZH2 variant | R25Q | K629E | A103V | E640K | I646F | R342W | V442D |
|  | D124V | V632A | A255T | E721Q | I708M | R46H | V621A |
|  | N130K | Y641N | A500D | E740A | I708T | R497P | V621M |
|  | P132L | Y641F | A576V | E740G | K210E | R497Q | V657A |
|  | P132S | Y641S | A651D | E740K | K243Q | R504G | V674L |
|  | Y133C | Y641H | A651P | E740Q | K491R | R504T | V674M |
|  | M134T | Y641C | A651V | F145L | K510R | R561H | V675M |
|  | F145L | V674M | A677T | F665L | K563E | R561L | V737D |
|  | E148G | A677G | A677V | G135R | K569E | R561P | Y133H |
|  | E148K | A687V | A687V | G155R | K680M | R578Q | Y133N |
|  | Y153C |  | C286Y | G159R | L128S | R608P | Y153C |
|  | Y153del |  | C289R | G159W | L149P | R653I | Y518P |
|  | K156E |  | C324R | G266E | L149R | R654I | Y641C |
|  | G159R |  | C536Y | G266R | L252V | R654K | Y641D |
|  | D233V |  | C543R | G5R | L26P | R679C | Y641N |
|  | H279R |  | C543Y | G623V | L666V | R679H | Y658C |
|  | H279Y |  | C553S | G643E | L669F | R679L | Y661N |
|  | K563E |  | C560Y | G643R | L669S | R679P | Y726C |
|  | H613Y |  | C566R | G643V | L669V | R679S | Y726D |
|  | V621M |  | C566S | G655A | M121I | R685C | Y726H |
|  | K634E |  | C566Y | G655E | N102S | R685G | Y728C |
|  | S647C |  | C585S | G655R | N182K | R685H | Y728H |
|  | D649G |  | C601Y | G681R | N317S | R727I |  |
|  | A651T |  | C642R | G704S | N555K | S280C |  |
|  | Y658N |  | D136G | G709V | N668Y | S474Y |  |
|  | D659V |  | D142Y | H129Q | N670K | S600R |  |
|  | S664N |  | D185H | H129R | N688K | S610A |  |
|  | N668Y |  | D192N | H279Y | N688T | S664G |  |
|  | A677T |  | D293G | H282Q | N688Y | S664N |  |
|  | R679C |  | D511E | H297Y | P132R | S664R |  |
|  | R679H |  | D652G | H525N | P132T | S690A |  |
|  | H689Y |  | D659A | H689R | P558L | S690L |  |
|  | S690L |  | D659E | H689Y | P572L | S690P |  |
|  | H706R |  | D659G | H706R | P693L | S729I |  |
|  | Y728* |  | D659H | I131F | Q648H | T261N |  |
|  | S729G |  | D659V | I131L | Q648K | T388M |  |
|  | Q730H |  | D725Y | I131N | R16Q | T568I |  |
|  | Y736C |  | E125K | I131S | R25Q | T678I |  |
|  | E740K |  | E173Q | I146T | R288Q | T718S |  |
|  | E740V |  | E249K | I55M | R342Q | V13A |  |

Taken together, this protein-level analysis of the mutational spectrum of WS strongly indicates that the associated pathogenic EZH2 variants are non-loss-of-function, and potentially dominant negative in nature.

### Weaver syndrome-associated EZH2 variants have dominant negative effects on PRC2 activity

To functionally evaluate whether WS-associated EZH2 variants are indeed dominant negative, we sought to compare the effects of heterozygous WS-variant expression versus the *Ezh2* heterozygous null condition. To this end, we generated *Ezh2* heterozygous knockout mESCs (Supplemental Fig. S1) and then exogenously expressed FLAG/HA-tagged human wild-type EZH2 (EZH2-WT) or variant EZH2 (Fig. 2A). We chose to investigate 10 EZH2 variants which are broadly reflective of the genetic and phenotypic heterogeneity of WS (Fig. 1A, Table 1). For comparison, we also expressed the well-characterised lymphoma-associated EZH2-Y641F gain-of-function variant (Supplemental Fig. S2), which is known to increase H3K27me3 and decrease H3K27me2 when expressed in cells (Sneeringer et al. 2010; McCabe et al. 2012; Conway et al. 2015). The exogenous WT and variant EZH2 proteins were expressed at equal levels.

**Figure 2.**
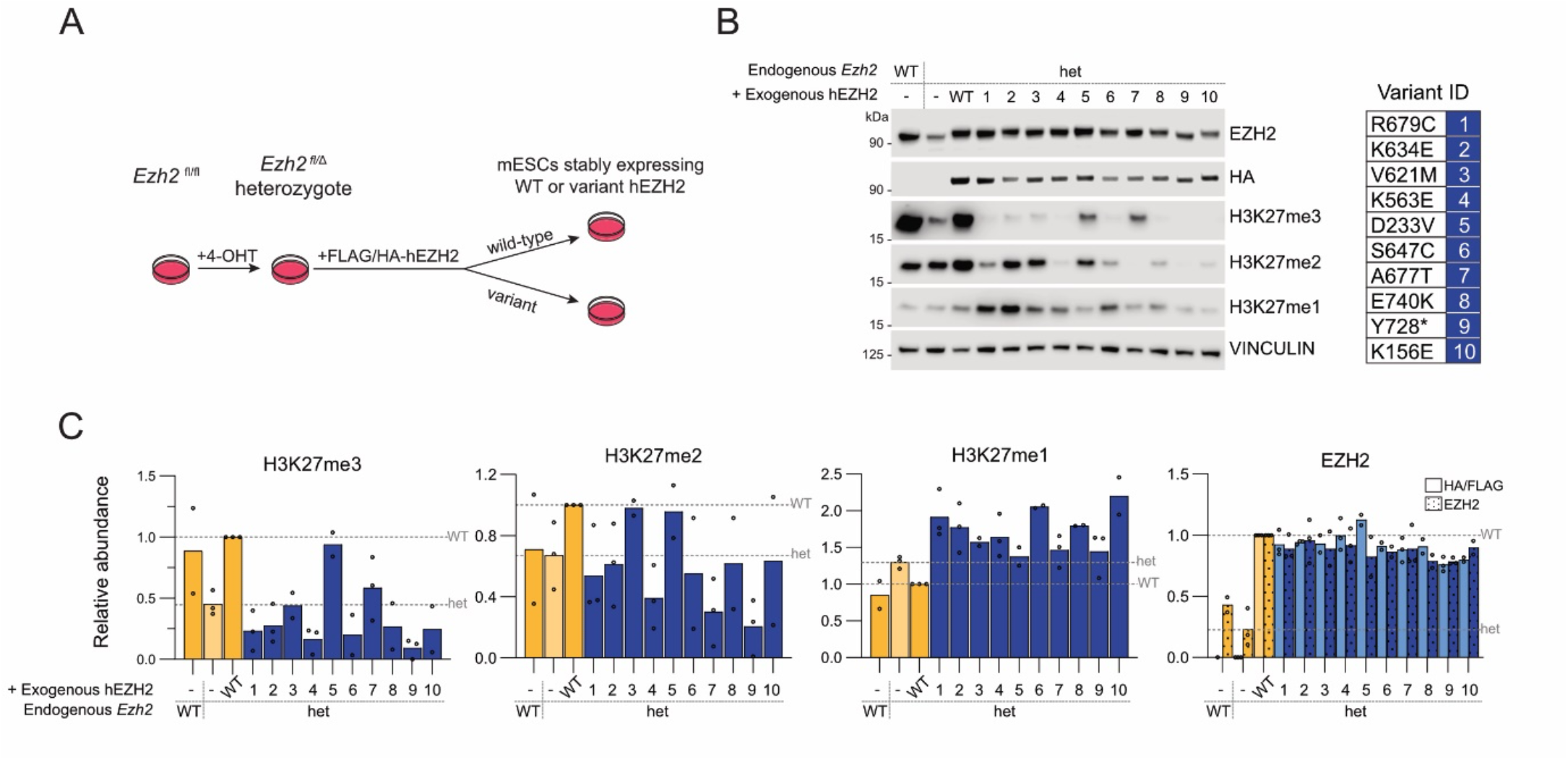
Weaver syndrome-associated EZH2 variants dominant negatively impair global PRC2 activity. **A**. Schematic of the isogenic stem cell model system used to generate the data presented in panels (b) and (c), and for all subsequent figures. **B**. Representative chemiluminescent immunoblots for HA, EZH2 and H3K27 modifications states, probing whole cell lysates from the indicated cell lines. VINCULIN is shown as a loading control. **C**. Relative quantification of fluorescent immunoblot analyses of EZH2 and the HA/FLAG epitope tag, H3K27me3, H3K27me2, and H3K27me1 abundance in the indicated cell lines. Signal intensity values were normalized to a loading control and then expressed relative to the *Ezh2*^fl/Δ^ + hEZH2-WT cell line. The dashed lines represent the normalised *Ezh2*^*fl*/Δ^ (het) and *Ezh2*^*fl*/Δ^ + hEZH2-WT (WT) values. Data values represent n=2 or n=3 independent biological replicates whose mean values are plotted as bars. *Abbreviations: het, heterozygote; WT, wild-type; 4-OHT, 4-hydroxytamoxifen*.

When expressed in *Ezh1/2* double knockout mESCs, which lack all H3K27 methylation (Højfeldt et al. 2018), all 10 WS-associated variants possessed some degree of H3K27 methyltransferase activity (Supplemental Fig. S2A-C), supporting their partial but not total loss of function. However, we did not find evidence supporting additive effects on PRC2 activity for these variants when expressed in the *Ezh2*^fl/Δ^ heterozygote (Fig. 2B-C). Rather, we observed apparent negative interference on the activity of EZH2 expressed from the endogenous WT allele. Immunoblot analyses for H3K27 methylation showed that, compared to cells exogenously expressing EZH2-WT, 9 out of the 10 WS-associated EZH2 variants induced global reductions in H3K27me3 and/or H3K27me2 (Fig. 2B-C). Importantly, these reductions occurred in a dominant-negative manner since the average decrease was more severe versus the parental *Ezh2*^fl/Δ^ heterozygote knockout cell line (Fig. 2B-C). Interestingly, EZH2-A677T [variant 7] only exhibited dominant-negative interference for H3K27me2. Moreover, on closer inspection of the original reporting article, the outlier EZH2-D233V variant (variant 5) that had WT-comparable H3K27me3 and H3K27me2 levels was associated with a relatively mild clinical presentation of WS (Tatton-Brown et al. 2013), supporting the utility of this cellular assay as a functional readout of variant severity. For the other 9 WS-associated EZH2 variants studied, the strong global reductions in H3K27 methylation were associated with 2-to-3-fold increases in global H3K27ac (Supplemental Fig. S2D-E). Collectively, these data comprise the first experimental evidence supporting dominant negative effects on global H3K27me3 and/or H3K27me2 levels by WS-associated EZH2 variants.

### Weaver syndrome-associated EZH2 variants deplete intergenic H3K27 methylations and impair global chromatin compaction

We previously speculated that disruption of intergenic H3K27me2/3 could contribute to the aetiology of Weaver syndrome (Deevy and Bracken 2019). To test this, we performed exogenous reference genome-normalised ChIP-seq (ChIP-Rx) for H3K27me2, H3K27me3, and H3K27ac. These and all subsequent analyses focus on a subset of 5 WS-associated EZH2 variants (Fig. 3A, Supplemental Fig. S3A-B) that were selected based on their frequency of occurrence, striking clinical phenotype, or both: R679C (variant 1), K634E (variant 2), K563E (variant 4), A677T (variant 7) and Y728* (variant 9).

**Figure 3.**
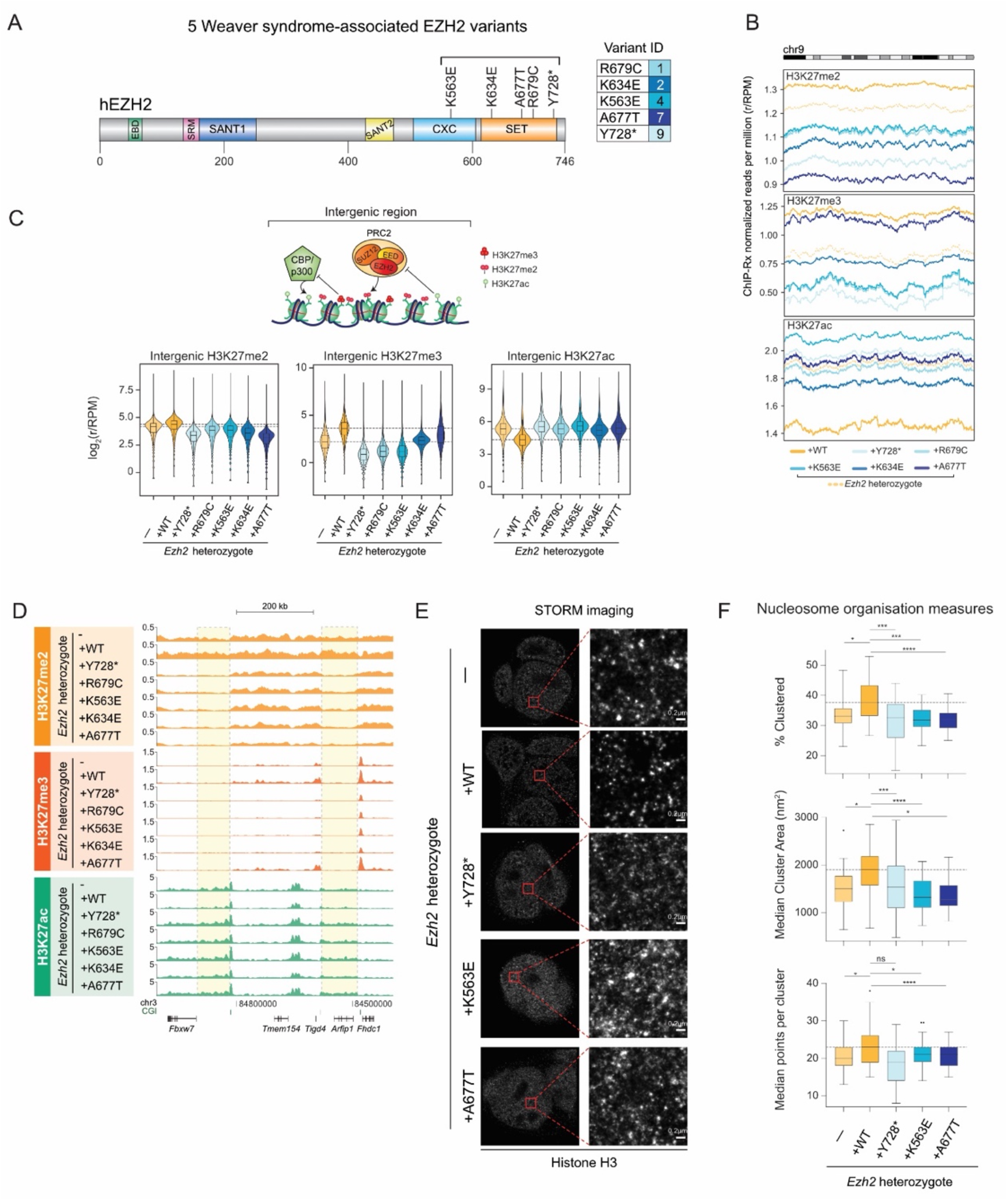
Weaver syndrome-associated EZH2 variants deplete intergenic H3K27 methylations and impair global chromatin compaction. **A**. Schematic of the human EZH2 protein (isoform 1; Q15910-1) highlighting the five Weaver syndrome-associated variants selected for further study. **B**. Rolling average plots presenting the ChIP-Rx normalized reads per million for H3K27me2, H3K27me3 and H3K27ac ChIP-Rx signal in cells expressing WS EZH2 variants compared to EZH2-WT across the whole of chromosome 9 (272477 10kb bins). **C**. (Top) Schematic illustrating the predominant H3K27 methylation states mediated by PRC2 at Polycomb target genes (left; H3K27me3) versus intergenic regions (right; H3K27me2). Also depicted is the antagonistic relationship between PRC2-mediated H3K27me2/3 and CPB/p300-mediated H3K27ac. (Bottom) Violin plot representations of the log_2_ ChIP-Rx normalized reads per million of H3K27me2, H3K27me3, and H3K27ac enrichments in the indicated cell lines across 188,538 10kb bins representing all intergenic regions. The dashed lines represent the EZH2-WT rescue cell line values. **D**. UCSC genome browser tracks showing ChIP-Rx normalised reads of H3K27me2, H3K27me3 and H3K27ac at a ∼600 kb genomic locus in the indicated cell lines. This locus includes a repressed PRC2 target gene (*Fhdc1*), two active genes (*Arfip1, Fbxw7*) and intervening non-genic regions. Representative gene body and intergenic regions are highlighted. CGI = CpG island. **E**. Representative stochastic optical reconstruction microscopy (STORM) images of the indicated cell lines stained for Histone 3. **F**. STORM quantifications of percentage clustered (top), median cluster area (middle), and median points per cluster (bottom) of H3. Data are represented as mean ± standard deviation. *Abbreviations: ChIP-Rx*, exogenous reference genome-normalised ChIP-seq; *EBD, EED-binding domain; SRM, stimulatory response motif*.; r/RPM, ChIP-Rx normalized reads per million; *STORM, stochastic optical reconstruction microscopy; WT, wild-type*.

Consistent with our immunoblot analyses (Fig. 2B-C), we found that all 5 chosen WS-associated EZH2 variants exhibited reduced H3K27me2 and H3K27me3 genome wide compared to EZH2-WT (Supplemental Fig. S3C), as shown along the length of chromosome 9 as an example (Fig. 3B). Specific examination of intergenic regions confirmed that all 5 WS-associated EZH2 variants depleted intergenic H3K27me2 and H3K27me3 compared to EZH2-WT (Fig. 3C). The same relative trend was observed for H3K27me2 and H3K27me3 at gene bodies (Supplemental Fig. S3D). The 5 WS-associated variants similarly exhibited lower intergenic (Fig. 3C) and gene-body (Supplemental Fig. S3D) H3K27me2 compared to the parental *Ezh2*^fl/Δ^ heterozygote. Except for EZH2-A677T, the WS-associated EZH2 variants also displayed reduced intergenic (Fig. 3C) and gene-body (Supplemental Fig. S3D) H3K27me3 compared to the parental *Ezh2*^fl/Δ^ heterozygote. This observation further highlights the potential of WS-associated EZH2 variants to exert dominant-negative effects on PRC2 activity.

Consistent again with our immunoblot analyses (Supplemental Fig. S2D-E), all 5 WS-associated EZH2 mutations increased genome-wide, intergenic, and gene-body H3K27ac (Fig. 3B-C, Supplemental Fig. S3C-D). In cells expressing these WS-associated EZH2 variants, H3K27ac tended to be gained on chromatin wherever H3K27me3 or H3K27me2 were lost, suggesting that the increase in H3K27ac may be the direct result of an increased availability of unmethylated H3K27 substrate (Fig. 3D; Supplemental Fig. S3E). Given the scale of the changes we observed on the landscape of H3K27 modification in response to WS-associated EZH2 variants, we next sought to investigate whether these changes correspond to an altered chromatin architecture. To do this, we performed H3 staining followed by stochastic optical reconstruction microscopy (STORM) imaging to assess chromatin architecture via the distribution of nucleosomes into clusters (Ricci et al. 2015). Relative to EZH2-WT, cells expressing WS-associated EZH2 variants had less compact chromatin with H3 present in fewer and less densely packed nucleosome clusters, as represented visually by the more diffuse histone H3 signal in Fig. 3E, and quantified in Fig. 3F. These data demonstrate that chromatin is more densely packed in cells expressing EZH2-WT compared to cells expressing WS-associated EZH2 variants. Taken together, the ChIP-Rx and STORM data point to a role for PRC2 in regulating global chromatin compaction and implicate its impairment in the aetiology of Weaver syndrome.

### Weaver syndrome-associated EZH2 variants differ in their effects on H3K27me3 at Polycomb target genes

We next examined the consequences of WS-associated EZH2 variants on the ability of PRC2 to deposit H3K27me3 at Polycomb target genes (Fig. 4A). 4 of the 5 WS-associated EZH2 variants had reduced mean levels of H3K27me3 at Polycomb target gene promoters (Fig. 4B). Surprisingly, the EZH2-A677T variant instead showed moderate increases in mean H3K27me3 deposition at these loci (Fig. 4B-C). We observed converse changes in mean H3K27me2 levels at Polycomb target promoters: 4 out of the 5 EZH2 variants showed increased levels of promoter H3K27me2, whereas the EZH2-A677T variant depleted H3K27me2 from these sites (Fig. 4B-C). We noted that the pattern of increased H3K27me3 and reduced H3K27me2 at Polycomb target genes caused by the EZH2-A677T variant was similar to, albeit less profound than, the oncogenic gain-of-function EZH2-Y641F and EZH2-A677G variants reported in lymphoma to have enhanced ability to convert H3K27me2 to H3K27me3 as a result of their enlarged lysine tunnels (Supplemental Fig. S4A-B) (McCabe et al. 2012). Supporting this, structural modelling of in silico mutated EZH2 points to an analogous mutational effect potentially underpinning the similarities between EZH2-A677T and EZH2-Y641F (Supplemental Fig. S4C). Specifically, the variant A677T residue is expected to cause steric clashes with Y641 (red discs in Supplemental Fig. S4C, centre panel). This could in turn shift Y641 and therefore enlarge the lysine tunnel, as predicted to occur in lymphoma when (1) Y641 loses its hydroxyl upon mutation to phenylalanine in the EZH2-Y641F variant (Supplemental Fig. S4D, right panel), and (2) A677 is mutated to a smaller glycine residue in the EZH2-A677G variant (McCabe et al. 2012). Taken together, these data indicate that although most (4 out of 5 tested) WS-associated EZH2 variants reduce the ability of PRC2 to deposit H3K27me3 at Polycomb target genes, this will not necessarily hold true for all pathogenic variants.

**Figure 4.**
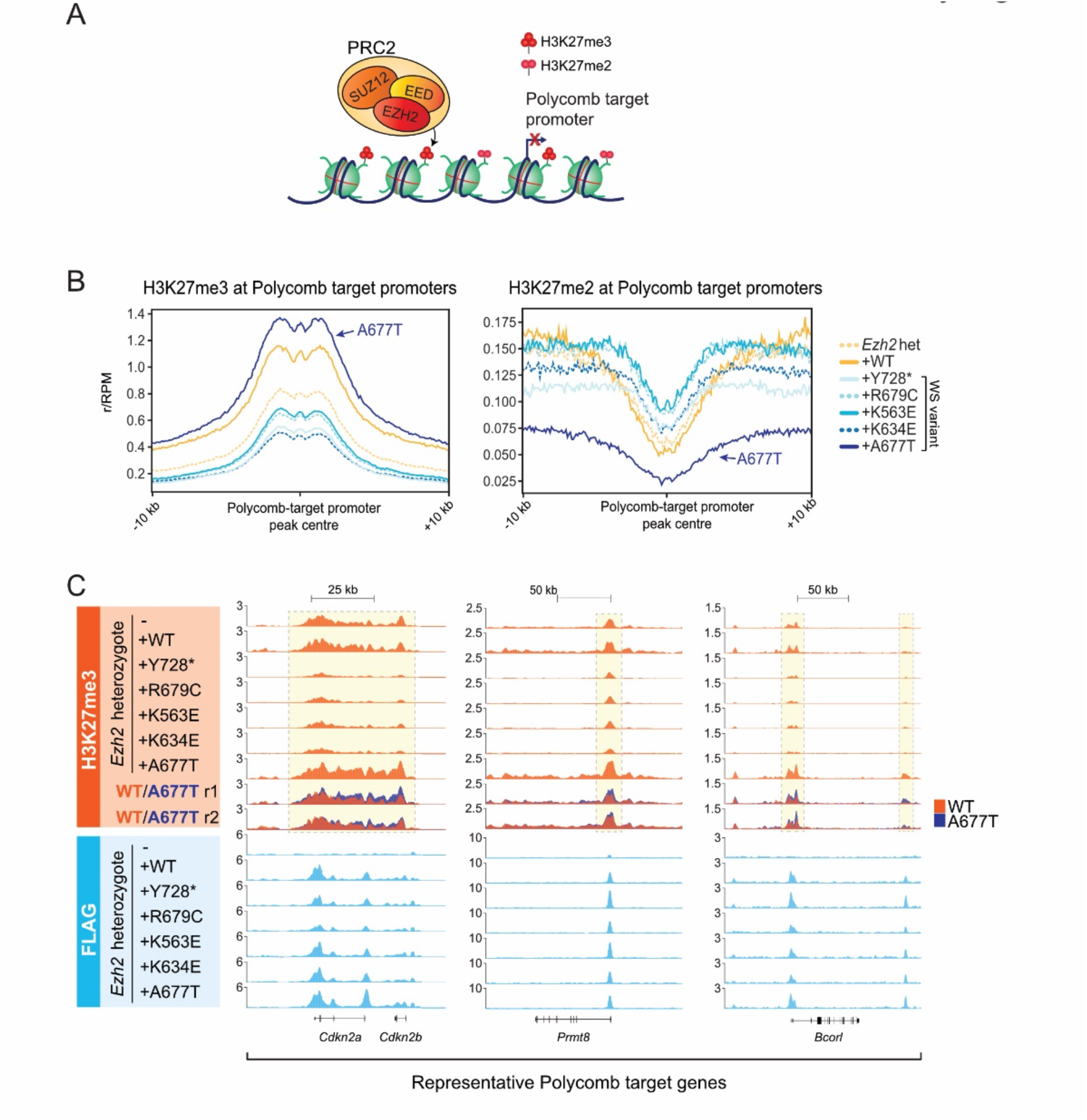
Weaver syndrome-associated EZH2 variants differ in their effects on H3K27me3 at Polycomb target genes. **A**. Schematic illustrating PRC2-mediated H3K27me3/me2 deposition at transcriptionally repressed Polycomb target promoters. **B**. Average ChIP-Rx signal profiles for H3K27me3 (left) and H3K27me2 (right) at Polycomb target promoters (n = 3870) in the indicated cell lines. Control cell lines are coloured in yellows while cells expressing Weaver syndrome EZH2 variants are coloured in blues. **C**. UCSC genome browser tracks showing ChIP-Rx normalised reads of H3K27me3 (top) and FLAG (bottom) at the indicated Polycomb target gene loci (*Prmt8, Bcorl, Cdkn2a/b*) in *Ezh2*^*fl/Δ*^ mESCs with or without stable exogenous expression of indicated FLAG-tagged EZH2 constructs. Also shown are overlayed tracks from two biologically independent ChIP-Rx experiments, directly comparing the H3K27me3 enrichment from the cell lines expressing either EZH2-WT (red) or EZH2-A677T (blue). *Abbreviations: ChIP-Rx*, exogenous reference genome-normalised ChIP-seq; *het, heterozygote; r1, replicate 1; r2, replicate 2; WT, wild type*.

### The growth restriction-associated EZH2-A733T variant induces opposing global H3K27 modification changes

A heterozygous gain-of-function mutation in EZH2 is implicated in an opposing developmental phenotype characterized by growth restriction, and the resulting EZH2 variant (p.A733T; Fig. 5A) demonstrates increased HMT activity in vitro (Choufani et al. 2020). We wished to determine the consequences of this gain-of-function EZH2 variant on the genome-wide deposition of H3K27me2 and H3K27me3 compared to WS-associated EZH2 variants. To this end, we expressed either EZH2-WT, the growth restriction-associated EZH2-A733T variant, or the WS-associated EZH2-Y728* variant in *Ezh2*^fl/Δ^ heterozygous mESCs and performed immunoblot analyses (Fig. 5B). As expected, this showed that the EZH2-A733T variant induced stark global increases in H3K27me3, coupled with global reductions in H3K27me1, H3K27me2, and H3K27ac (Fig. 5B). Next, we performed ChIP-Rx of H3K27me2, H3K27me3, and H3K27ac on *Ezh2*^fl/Δ^ heterozygous mESCs expressing either WT EZH2, the growth-restriction-associated EZH2-A733T variant, or 1 of 2 representative WS-associated variants: EZH2-Y728* or EZH2-A677T (Supplemental Fig. S5A-B). In direct contrast to the WS-associated EZH2 variants, the growth restriction-associated EZH2-A733T variant caused genome-wide and intergenic increases of H3K27me3 coupled with decreased H3K27ac (Fig. 5C and Supplemental Fig. S5C-D). We observed high levels of H3K27me3 spreading pervasively from PRC2 bound sites into surrounding intergenic regions, with reciprocal H3K27ac reductions (Fig. 5D). Accordingly, a genome-wide correlation analysis showed that H3K27ac was reduced where H3K27me3 was gained (Fig. 5E, right panel). H3K27me2 was also reduced in these same regions (Fig. 5E, left panel) as it was converted to H3K27me3 (Supplemental Fig. S5D). The expression of EZH2-A733T also increased mean levels of H3K27me3 at PRC2 target gene promoters with a corresponding decrease in mean H3K27me2 (Fig. 6A) – consistent with its gain-of-function mechanism (Supplemental Fig. S5E).

**Figure 5.**
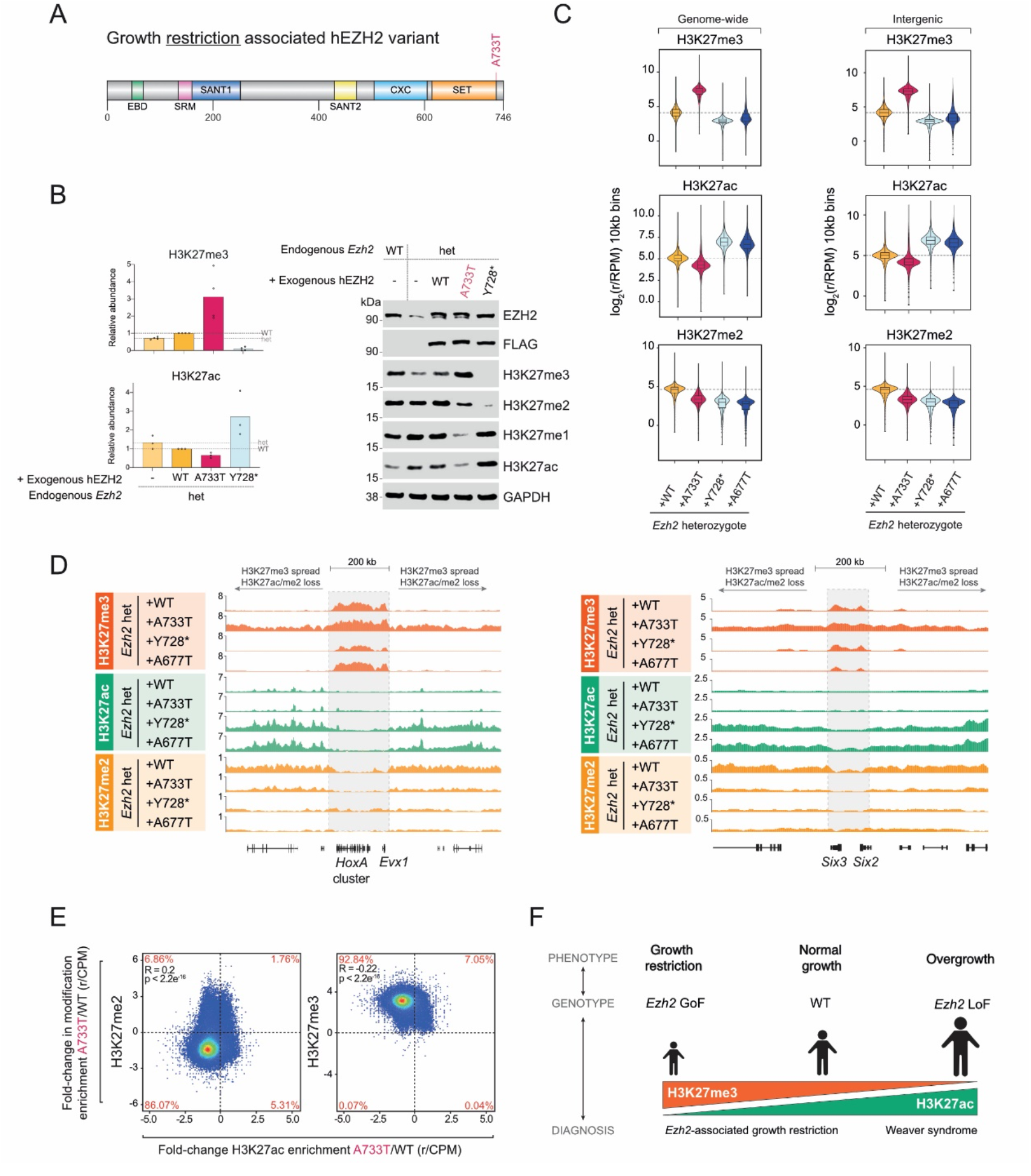
The growth restriction-associated EZH2-A733T variant induces opposing global H3K27 modification changes. **A**. Schematic of the human EZH2 protein showing the location of the growth restriction-associated A733T variant. **B**. (Left) Relative quantification of immunoblot analyses of H3K27me3 (top) and H3K27ac (bottom) abundance in the indicated cell lines. Yellows represent the control cell lines, pink represents the growth restriction-associated variant, and blue represents the Weaver syndrome-associated variant. Dashed lines represent the normalised *Ezh2*^*fl/Δ*^ (het) and *Ezh2*^*fl/Δ*^ + EZH2-WT (WT) values. Data values represent independent biological replicates, and their mean values are plotted as bars. (Right) Representative immunoblots for FLAG, EZH2, and H3K27 modification states, probing whole cell lysates from the indicated cell lines. GAPDH is shown as a loading control. **C**. Violin plot representations of the log2 ChIP-Rx normalized reads per million of H3K27me3, H3K27ac and H3K27me2 enrichments in the indicated cell lines genome-wide (left) and across 188,538 10kb bins representing all intergenic regions (right). Dashed lines represent the *Ezh2*^*fl/Δ*^ + EZH2-WT values. **D**. UCSC genome browser tracks showing ChIP-Rx normalised reads of H3K27me3, H3K27ac and H3K27me2 at the *HoxA* gene cluster (left) and *Six2/3* genes (right) in the indicated cell lines. Note how the EZH2-A733T growth restriction variant causes aberrant spread of H3K27me3 left and right of these loci at the expense of H3K27me2 and H3K27ac. **E**. Correlation plots demonstrating the relationship between changes in genome-wide ChIP-Rx enrichment of H3K27ac and H3K27me2 (left) or H3K27me3 (right) in the EZH2-A733T growth restriction variant compared to EZH2-WT. The percentage of 10kb bins in each quadrant is indicated. The p-value and correlation coefficient were calculated using Pearson’s correlation. **F**. Schematic illustrating a proposed model for the roles of H3K27 methylation and acetylation levels in controlling human growth during development, where growth restriction associates with hypermethylation and hypoacetylation whereas overgrowth associates with hyperacetylation and hypomethylation. The balance between histone H3K27 methylation and acetylation is important for maintaining normal growth. *Abbreviations: ChIP-Rx, exogenous reference genome-normalised ChIP-seq; het, heterozygote; r/RPM, ChIP-Rx normalized reads per million; WT, wild type*.

Taken together, these data suggest that global H3K27 hyper-trimethylation and H3K27 hypo-acetylation are characteristic of the growth restriction-associated EZH2-A733T variant, and vice versa for WS overgrowth-associated EZH2 variants (Fig. 5F), lending support to their potential biological relevance in the opposing developmental phenotypes.

## Discussion

Here we present an investigation of the molecular effects of a broad range of WS-associated EZH2 variants that cause developmental overgrowth, as well as an opposing EZH2 variant associated with developmental growth restriction. We provide the first evidence that WS-associated EZH2 mutations can confer dominant negative effects on the global H3K27 methylation activities of PRC2. Strikingly, we observed directly opposing effects for an EZH2 variant associated with developmental growth restriction. We propose that these opposing chromatin defects are central to the underlying pathophysiology of EZH2-related growth disorders.

### Weaver syndrome-associated EZH2 variants can dominant-negatively affect PRC2 activity

Here, we used 3D protein structural data to predict that WS-associated EZH2 variants are non-loss-of-function, with likely dominant-negative effects. This prediction is consistent with the known mutational spectrum and inheritance pattern of WS. Previously published models of WS suggest that the associated EZH2 variants are partial loss-of-function, but they did not include a matched comparison against the EZH2 heterozygous condition to allow testing for dominant interference (Cohen et al. 2016; Lui et al. 2018; Jani et al. 2019; Choufani et al. 2020; Gao et al. 2024). We experimentally addressed this question by creating an *Ezh2* heterozygous knockout mESC line in which we exogenously expressed human WS-associated EZH2 variants (Fig. 2A), noting that the mouse (Q61188) and human (Q15910) EZH2 proteins share 98.4% amino acid sequence identity. In doing this, we were able to show for the first time that PRC2 activity tends to be more severely impaired by the heterozygous expression of WS-associated EZH2 variants compared to the isogenic heterozygous loss of EZH2.

We hypothesize that these dominant-negative effects may be mediated through the heterozygous expression of hypomorphic WS-associated EZH2 variants that occupy normal PRC2 binding sites but deposit less H3K27me3/me2. We show that the expression of WS-associated EZH2 variants preserves the locations but reduces the levels of both H3K27me3 and H3K27me2. H3K27me3 is known to allosterically activate PRC2 and stabilize its binding on chromatin, while PRC2 was recently shown to have binding affinity for both H3K27me2 H3K27me3 modified nucleosomes (Lee et al. 2018; Lukauskas et al. 2024). As such, allosteric activation and binding of PRC2 is expected to decreased as a result of reduced H3K27me3/me2, irrespective of whether it contains WT or variant EZH2. Therefore, we propose a dominant negative mechanism in which WS-associated EZH2 variants impair both the allosteric activation and binding of WT EZH2 containing PRC2.

### A dominant negative mutational mechanism may explain the lack of early truncating mutations in Weaver syndrome

Our discovery of dominant-negative effects by WS-associated EZH2 variants may help answer the question posed by Lui et al. (2018): if the causative mutations are indeed loss-of-function, why have no early truncating mutations of EZH2 been reported in WS? One possibility proposed by Gao et al. (2024) is that manifestation of the Weaver phenotype may require biallelic EZH2 protein expression, irrespective of catalytic activity. Our data agree that normal EZH2 protein levels may be necessary but suggest that perturbed catalytic activity is central to the phenotype. We rather propose that the Weaver phenotype is in most cases a consequence of dominant-negative interference of PRC2 activity during development, which necessitates the expression of both a WT and pathogenic protein product.

Another potential explanation is that heterozygous-null mutations in EZH2 are lethal. However, as of 20th March 2024, the Database of Chromosomal Imbalance and Phenotype in Humans Using Ensembl Resources (DECIPHER) reports 94 patients with heterozygous deletions encompassing EZH2 (Firth et al. 2009), so this is not likely. A third possibility is that heterozygous-null mutations in EZH2 cause a phenotype that is distinct from Weaver syndrome. Indeed, among the subset of 82 heterozygous whole-gene deletion EZH2 variants with phenotype documentation available in DECIPHER, none are associated with overgrowth or tall stature. On the contrary, short stature and small for gestational age are, respectively, the second and third most common phenotypes reported.

Finally, it is possible that heterozygous-null mutations in EZH2 are clinically silent. Supporting this, several early truncating variants in EZH2 have been uploaded to ClinVar as of March 2024 (Landrum et al. 2018). In this scenario, EZH2 haploinsufficiency may not sufficiently impair PRC2 activity to produce the Weaver phenotype, which we speculate may be due to partial compensation by EZH1. Although EZH1 catalytic activity and abundance in ESCs is substantially lower than EZH2 (Margueron et al. 2008; Shen et al. 2008; Streubel et al. 2018), it might sufficiently compensate for EZH2 at later stages of development when its expression levels increase. Interestingly, the mutational spectrum for the closely related SUZ12-mutant Imagawa Matsumoto syndrome includes heterozygous early truncating variants, whole-gene deletion variants, and even homozygous missense variants (Cyrus et al. 2019b; Park and Jang 2023; Yuksel Ulker et al. 2023). Since loss of SUZ12 function would affect both EZH1- and EZH2-containing PRC2 complexes, the contrasting mutational spectrum of Imagawa Matsumoto versus Weaver syndrome supports functional compensation for heterozygous-null mutations in EZH2 by EZH1.

Overall, our findings agree with prior reports that WS-associated EZH2 variants are partial loss-of-function and extend them by providing the first evidence supporting their dominant negative effects.

### What do we learn about the link between Weaver syndrome-associated EZH2 variants and cancer?

Individuals with WS are highly predisposed to developing cancers (Kamien et al. 2018). Indeed, some WS variants are directly found in cancer malignancies, including EZH2-P132L, EZH2-R679C (variant 1), EZH2-E740K (variant 8), and EZH2-Y728* (variant 9). Usemann et al. (2016) found the heterozygous EZH2-P132L variant in a young patient with WS who developed acute myeloid leukemia. EZH2-R679C was detected in the homozygous condition in a secondary myelofibrosis case, while EZH2-Y728* was found to remain in the heterozygous condition in a case of primary myelofibrosis that transformed into acute myeloid leukemia (Ernst et al. 2010, table S3). Nevejan et al. (2020) identified EZH2-E740K as the driver mutation in a case of donor cell-derived acute myeloid leukemia after stem cell transplantation. These observations provide potentially relevant insight for predicting cancer predispositions in affected WS cases. There are also documented examples of alternative variants occurring in cancer at the same residues mutated in WS, most notably including the oncogenic EZH2-A677G variant (McCabe et al. 2012; Gu 2016), which shares the same residue as the WS-associated EZH2-A677T characterised in this study.

Here, we show the WS-associated EZH2-A677T variant behaves somewhat like gain-of-function EZH2-Y641F and EZH2-A677G variants previously reported in B-cell lymphoma (Sneeringer et al. 2010; McCabe et al. 2012). Like the B-cell lymphoma variants, we show that EZH2-A677T has an enhanced ability to convert H3K27me2 to H3K27me3. In accordance with this “oncogene-like” mechanism of action, the individual with WS caused by this EZH2-A677T variant developed both neuroblastoma and acute lymphoblastic leukemia at just 13 months-old (Tatton-Brown et al. 2013). Considering these potentially cancer-relevant insights, we propose that genomic profiling of H3K27 modifications upon ectopic expression of WS-associated EZH2 variants may represent a useful means to predict cancer predisposition in individuals.

### Assessing genotype-phenotype correlations for chromatinopathies using our experimental framework

We show that the degree of dominant-negative impairment of PRC2 functions by WS-associated EZH2 variants correlates with the clinical phenotype. For example, we identified one EZH2 variant reported to cause familial Weaver syndrome (EZH2-D233V) that did not impair the global histone methylation activities of PRC2. In line with this, we noted that the individuals diagnosed with WS who had this mutation had relatively mild clinical presentations, with heights no more than +2 standard deviations from the norm (SD), and occipito-frontal circumferences no more than +0.6 SD (Tatton-Brown and Rahman 2013). In contrast, the WS-associated EZH2 variant Y728* was the most severely impaired in our cellular assays of HMT activity, and accordingly, the individual with this variant had a strong presentation of clinical overgrowth, with a recorded height +7.6 SD and occipito-frontal circumference +2.7 SD (Tatton-Brown et al. 2011).

Therefore, we propose that the cellular assay and investigative strategy designed here could be extended to study other growth disorders caused by mutations in chromatin regulators, such as SUZ12, EED, ASXL1-3, NSD1, DNMT3a, CHD8, ARID1B, BRM, and CBP/EP300 (Tatton-Brown et al. 2017; Alfert et al. 2019; Gamu and Gibson 2020; Cuddapah et al. 2021). Notably, loss-of-function mutations in CBP/P300 H3K27 acetyltransferases cause Rubinstein-Taybi syndrome, which is characterized by short stature, microcephaly and other features directly opposing WS clinical presentation (Gamu and Gibson 2020). It is possible that the defects in H3K27 acetyltransferase activity could lead to lower levels of H3K27ac across the genome, similar to the effects of the growth restriction-associated EZH2 variant studied here, and in contrast to the elevated levels of H3K27ac observed in our cells expressing Weaver syndrome-associated variants.

In summary, we have shown here that cellular modelling of mutations to identify common changes across overgrowth syndromes, and directly opposing changes in growth restriction models, is a powerful way to pinpoint cellular defects that are central to these disorders.

## Materials and Methods

### Cell Culture and cell lines

Mouse embryonic stem cells (ESCs) were grown on 0.1% gelatin-coated culture dishes in Glasgow Minimum Essential Medium (Sigma) supplemented with 20% (v/v) heat-inactivated FBS (Gibco), 100 U/mL Penicillin and 100 μg/mL Streptomycin (Gibco), 50 μM β-mercaptoethanol, 1:100 GlutaMAX (Gibco), 1:100 non-essential amino acids (Gibco), 1 mM sodium pyruvate (Gibco), 1:500 leukaemia inhibitory factor (LIF; produced in-house), 3 μM CHIR99021 (Cayman Chemical Company) and 1 μM PD0325901 (Cayman Chemical Company). HEK293T cells (for lentivirus production) and NTERA-2 cells (for ChIP-Rx spike-in) were grown on TC-treated culture dishes in high glucose Dulbecco’s Modified Eagle’s Medium (Sigma) supplemented with 10% (v/v) FBS (Gibco), 100 U/mL Penicillin and 100 μg/mL Streptomycin (Gibco).

The heterozygous *Ezh2* knockout cell line was generated by treating the *Ezh2*^fl/fl^;Rosa26CreERT2 mESCs with 0.5 μM 4-OHT for 12 hours. Single cells were from this population were then sorted into 96-well culture plates using a FACSAria Fusion Flow Cytometer. Individual clones were screened at the gDNA level for the desired heterozygous deletion in *Ezh2*, and positive clonal populations were expanded for further validation at the mRNA and protein expression level.

Stable cell lines expressing exogenous hEZH2 constructs were generated by lentiviral transduction. A second-generation lentiviral system was used to produce lentivirus particles in HEK293T cultures. mESCs were seeded at a density of ∼2×104 cells per cm2 on day 0, then treated with virus on day 1 and day 2. Approximately 48 hours post-transduction, cells were washed with DPBS and treated with 1 μg/mL puromycin. Antibiotic selection was maintained for an empirically defined length of time (typically four to six days) to obtain putative stable cell lines, which were subsequently screened for expression of the intended transgene.

### Plasmids

Human EZH2 expression vectors were generated using Gateway cloning and the in-house pLENTI-EFIA-FLAG/HA lentiviral expression vector (Conway et al. 2018). Mutant hEZH2 ORFs were generated by Q5 Site-Directed Mutagenesis (NEB E0554S) as per the manufacturer’s protocol. Note that our hEZH2 ORF corresponds to EZH2 isoform 1 (Q15910-1, 746 amino acids).

### Immunoblotting

Whole-cell protein samples were prepared in ice cold High Salt buffer (50 mM Tris-HCl pH 7.2, 300 mM NaCl, 0.5% (v/v) NP-40, 1mM EDTA pH 7.4, 2 μg/mL aprotinin, 1 μg/mL leupeptin, 1mM PMSF). Cell suspensions were sonicated and then rotated at 4 °C for 20 minutes to ensure sufficient lysis. Lysates were next clarified by centrifugation at ≥ 20,000 RCF at 4°C for 20 minutes. Normalised protein lysates were denatured and separated on Bolt Bis-Tris (Invitrogen) or NuPAGE Bis-Tris (Invitrogen) gels, then transferred to nitrocellulose membranes (0.2 μM, Amersham). Membranes were blocked against nonspecific binding and then incubated with the relevant primary (overnight at 4 °C) and secondary (1h at RT) antibodies. Relative protein levels were determined by chemiluminescence or infrared fluorescence detection on a LI-COR Odyssey Fc. For details on the antibodies used, see Table 3.

**Table 3.** Antibodies used in this study. *Abbreviations: BD43, hybridoma line BD43; ChIP, chromatin immunoprecipitation*.

| Target | Source (identifier) | Western blot | ChIP |
| --- | --- | --- | --- |
| EZH2 | Bracken lab (BD43) | 1:10 | - |
| EZH2 | Abcam (ab191250) | 1:1000 | - |
| H3K27ac | Abcam (ab4729) | 1:1000 | 2µg |
| H3K27ac | Sigma-Aldrich (07-360) | 1:1000 | - |
| H3K27me1 | Active Motif (0321) | 1:2000 | - |
| H3K27me2 | Cell Signalling (9728S) | 1:2000 | 2.25µg |
| H3K27me3 | Cell Signalling (custom) | - | 1µg |
| H3K27me3 | Active Motif (0323) | 1:2500 | - |
| HA | Cell Signalling (3724) | 1:1000 | - |
| FLAG | Sigma-Aldrich (clone M2) | 1:1000 |  |
| ACTIN | Proteintech (60008-1-Ig) | 1:5000 | - |
| GAPDH | Proteintech (60004-1-Ig) | 1:5000 | - |
| VINCULIN | Cell Signalling (4650) | 1:2500 | - |
| Mouse IgG | Millipore (12-371) | - | 2µg |

### STORM imaging and analysis

For STORM imaging, 35m MMATTEK (P35G-1.5-14-C) plates were gelatinised with 0.1% gelatin. 500,000 mESC were seeded per plate for 20-22 h before 6 minutes fixation in 1:1 methanol: Ethanol at −20°C. Samples were blocked in blocking buffer (PBS with 10% donkey serum) for one hour at room temperature. Samples were incubated overnight at 4°C with H3 antibody (ab1791) 1:70 dilution in blocking buffer + 0.1% Triton X-100. Samples were incubated in 1:200 dilution of Alexafluor-647 secondary antibody in PBS in the dark for one hour at room temperature. Samples were then post-fixed in 2% PFA for 5 minutes before storing in PBS at 4°C until imaging. Direct STORM (dSTORM) imaging was performed using the Nikon N-STORM microscope equipped with a 1.49 NA CFI Apochromat TIRF objective, exciting the Alexa Fluor 647 dye with the 647 nm laser light in HILO (highly inclined and laminated optical sheet) mode (Tokunaga et al. 2008). The 405 nm laser light (activation laser) was used for reactivating the Alexa Fluor 647 into a fluorescent state. The activation laser power was automatically increased by the NIS software to keep the number of localizations per frame constant up to a maximum of 50% of the laser power. Each dSTORM acquisition consisted of 40 thousand images recorded with an Orca-Flash4.0 sCMOS camera (Hamamatsu) with an exposure time of 20 ms, a pixel size of 161.5 nm and a field of view of 128×128 pixels. During dSTORM acquisitions, cells were kept in imaging buffer (100 mM MEA, 1% glucose, 560 ug/mL Glucose Oxidase and 34 ug/mL Catalase in PBS). Two regions of interest (ROI) of 32×32 pixels for each dSTORM image were processed using ThunderSTORM (Ovesny et al. 2014), with a pre-detection wavelet filter (B-spline, scale 2, order 3), initial detection by non-maximum suppression (radius 1, threshold at one standard deviation of the F1 wavelet), and sub-pixel localization by integrated Gaussian point-spread function and maximum likelihood estimator with a fitting radius of 3 pixels. Detected localizations were filtered (intensity > 500 photons, sigma range of 50–500, and localization uncertainty < 20 nm). The filtered dataset was then corrected for sample drift (cross-correlation of images from five bins at a magnification of 5) and repeated localizations was removed by merging points which reappeared within 20 nm. STORM images were visualized using the normalized Gaussian method with a lateral uncertainty of 20 nm. Cluster analysis were performed thanks to a supervised machine learning approach using trained neural network (Williamson et al. 2020). The CAML (Cluster Analysis by Machine Learning) analysis workflow consisting of 3 stages and corresponding Python scripts, was used to: 1) prepare the data converting the x,y localization tables in a list of near-neighbor distances; 2) evaluate the input data with a trained model; 3) extract clustering information. The 87B144 model was used (Williamson et al. 2020), considering the possibility that more than 100 localizations per cluster could occur in our dataset.

### Quantitative ChIP

Cells were counted and normalized cell pellets (each of equal cell number, up to 50×106) were washed twice with DPBS before crosslinking for 10 minutes with 1% formaldehyde (Sigma) in DBPS. Crosslinking was quenched with 125 mM glycine for 5 minutes, followed by two PBS washes. Crosslinked cells were lysed in an appropriate volume of SDS-lysis buffer (100 mM NaCl, 50 mM Tris pH 8.1, 5 mM EDTA pH 8.0, 0.5% SDS, 2 μg/mL aprotinin, 1 μg/mL leupeptin, 1 mM PMSF). Chromatin was pelleted by centrifugation at 350 RCF for 5 minutes at room temperature. The supernatants were discarded, and chromatin resuspended in ChIP buffer (2:1 dilution of SDS-lysis buffer/Triton dilution buffer [100 mM Tris pH 8.6, 100 mM NaCl, 5 mM EDTA pH 8.0, 5% Triton X-100, 2 μg/mL aprotinin, 1 μg/mL leupeptin, 1 mM PMSF]). For quantitative ChIP analyses (ChIP-Rx), spike-in chromatin was prepared from NTERA-2 cells and added to mESC samples (1:10 cell number ratio) at this point in the preparation procedure.

Chromatin samples were sheared using a Branson Sfx150 Sonifier for a total sonication time of 4-5 min at 50% power and a duty cycle of 1 s on and 4 s off. ChIPs were performed overnight at 4 °C. For details on the antibodies used, see Table 3. Following overnight incubation, each sample received 40 μl of Protein G Dynabeads and was incubated for a further 3 h at 4 °C. Beads were washed three times in mixed micelle buffer (150 mM NaCl, 20 mM Tris pH 8.1, 5 mM EDTA pH 8.0, 5.2% Sucrose, 1% Triton X-100, 0.2% SDS), twice with buffer 500 (0.1% Sodium Deoxycholate, 1 mM EDTA pH 8.0, 50 mM HEPES pH7.5, 1% Triton X-100), twice with LiCl detergent wash (0.5% Sodium Deoxycholate, 1 mM EDTA pH 8.0, 250 mM LiCl, 0.5% NP-40, 10 mM Tris pH 8.0) and once wash with TE (10 mM Tris pH 8.0, 1 mM EDTA pH 8.0). Samples were eluted from beads using elution buffer (0.1 M NaHCO3, 1% SDS) while shaking for 1 h at 65 °C, then supernatants were incubated overnight at 65°C to reverse crosslinking. Enriched DNA fragments were Rnase (Thermo Fisher) and Proteinase K (Sigma) treated prior to purification using Qiagen MinElute PCR Purification Kit (Qiagen; 28006).

ChIP library preparation. ChIP-purified DNA was quantified by fluorimetry using the Qubit dsDNA High Sensitivity Assay Kit (Thermo Fisher), and 0.5–50 ng of DNA/ChIP was used for library preparation using the NEBNext Ultra II DNA Library Kit for Illumina (NEB E7645) and NEBNext Multiplex Oligos for Illumina (Index Primers Set 1 and 2; NEB E7335 and E7500). Library DNA was quantified using the Qubit, and size distributions were ascertained on a TapeStation (Agilent) using the D1000 ScreenTape assay reagents (Agilent; 5067-5583). This information was used to calculate pooling ratios for multiplex library sequencing. Pooled libraries were diluted and processed for either 75-bp single-end or 35-bp paired-end sequencing on an Illumina NextSeq 500 instrument (ID: NB501524) using the NextSeq 500 High Output v2 kit (75 cycles) (Illumina; FC-404-2005) in accordance with the manufacturer’s instructions.

### Computational predictions of the nature of EZH2 missense variants

FoldX 5.0 (Delgado et al. 2019)was run to estimate the change in Gibbs free energy of folding (ΔΔG) of missense mutations as the average of 3 replicates, using the EZH2 chain in the x-ray structure 5hyn (Justin et al. 2016). The FoldX “RepairPDB” command was run before modelling the mutations. The Extent of Disease Clustering (EDC) metric was calculated as previously described (Gerasimavicius et al. 2022). Leukemia-associated missense variants were obtained from COSMIC v99 (cancer.sanger.ac.uk).

### ChIP-Rx data analysis

Chromosome names for the human genome were modified with the prefix hg38_ and a metagenome was created by concatenating the mouse and human reference genomes (mm10 and hg38, respectively) before indexing with Bowtie 2 version 2.3.5 (Langmead and Salzberg 2012), as described (Orlando et al. 2014). Reads were aligned to the metagenome using Bowtie 2 with default parameters, with the exception of –no-unal. Non-unique read alignments were filtered out using SAMtools version 1.10 (Li et al. 2009) to exclude those with an alignment quality of <2, and the hg38_ prefix appended to chromosome names was used to separate reads as aligned to the reference mouse or spike-in human genomes. SAMtools was used to convert SAM files to BAM files and to remove duplicate aligned reads. Spike-in normalization factors were calculated for each ChIP using the formula for normalized reference-adjusted reads per million (RRPM) as described (Orlando et al. 2014) 65; 1 per million spike-in reads. BigWig files were generated using the bamCoverage tool from the deepTools suite version 3.5.0 (Ramírez et al. 2016) with a bin size of 10 and scaled using the RRPM normalization factor derived for each sample. BAM files were converted to BED format using the bamToBed function in BEDTools version 2.27.1 (Quinlan and Hall 2010) and used for peak calling with HOMER version 4.10 (Heinz et al. 2010), using BED files from input ChIP-Rx as a control. The computeMatrix function in deepTools was used to score the coverage of BigWigs in genomic regions of interest prior to generating average and tornado plots using the plotProfile and plotHeatmap functions respectively. The featureCounts program version 1.6.4 was used to calculate the average ChIP signal at the TSS of genes of interest from the BAM files. The average signals were multiplied by the spike-in normalisation factors and heatmaps were generated using the Graphpad software version 9.5.1.

The mm10 reference genome was split into 10-kb windows using the makewindows function in BEDTools, and regions in the ENCODE mm10 blacklist were removed using bedtools intersect with the -v and -wa flags (Amemiya et al. 2019). BEDTools coverage was used to summarize overlapping read alignments in BAM files within these bins and scaled using RRPM normalization. Resulting counts were used for the generation of chromosome-wide-coverage line graphs and XY density scatter plots. Promoter regions were identified by taking −1,500 bases upstream and +600 bases downstream of annotated transcription start sites in the Ensembl 98 mm10 genome annotation. Gene body regions were defined by taking the gene annotations as defined in the Ensembl 98 mm10 genome annotation and converted to the bed format using a custom script. The BEDTools complement tool was used to generate a BED file of intergenic regions by supplying the gene body BED file a FASTA index of the mm10 genome as input (generated using SAMtools faidx).

### In silico mutagenesis and structural modelling

In silico mutagenesis was carried out in the context of the structure of PRC2.2 with a substrate nucleosome (PDB 6WKR). In silico mutagenesis was carried out using PyMol and van der Waals clashes between the mutated amino acids to residues in their vicinity were visualised using the show_bumps script, with clashes represented by discs. Red discs represent severe clashes and their diameter increases with clashes severity. Green discs represent nearly clashes. In cases where more than one rotamer was possible, the rotamer that provides the least severe clashes is presented.

## Acknowledgements

We thank members of the Bracken lab for helpful discussions and critical reading of the manuscript. We are grateful to the Genomics Core at University College Dublin for expertise and help with next-generation sequencing. We thank Kristian Helin for providing the *Ezh2*^fl/fl^ and *Ezh1/2* dKO cell lines. Work in the Bracken lab was supported by the Irish Research Council Advanced Laureate Award (IRCLA/2019/21) and Science Foundation Ireland (SFI), under the SFI Investigators program (SFI/16/IA/4562), the Irish Cancer Society (CancersUnmetNeeds012) and the St. Vincent’s Foundation.

O.D. was supported by a PhD fellowship from the Irish Research Council Government of Ireland Postgraduate Scholarship Programme (GOIPG/2017/2009). E.T. was supported by a PhD fellowship from the Irish Research Council Government of Ireland Postgraduate Scholarship Programme (GOIPG/2019/3481). F.M. was supported by a PhD fellowship from Trinity College Dublin Provost Award. E.C was supported by an SFI/IRC Pathway Program grant (21/PATH-S/9384) and a Wellcome Trust Early Career Award [225152/Z/22/Z]. J.A.M. was supported by the European Research Council (PROT-STRUCT-DISEASE, grant no. 101001169). The work of the Pasini laboratory was supported by the Worldwide Cancer Research (22-0027); the Italian Association for Cancer Research, AIRC (IG-2017-20290 and IG 2022-27694); and by the European Research Council, ERC (EC-H2020-ERC-CoG-DissectPcG: 725268). This study makes use of data generated by the DECIPHER community. A full list of centres who contributed to the generation of the data is available from https://deciphergenomics.org/about/stats and via email from. DECIPHER is hosted by EMBL-EBI and funding for the DECIPHER project was provided by the Wellcome Trust [WT223718/Z/21/Z].

## Author Contributions

O.D. and A.P.B. conceived of the project and designed the experiments. O.D. engineered the EZH2-variant mESC cultures and performed most of the bench-based experiments. F.M. performed additional Western blot and ChIP-Rx experiments. C.M., F.M. and D.N. performed the bioinformatics analyses.

E.C. and S.R. performed STORM imaging experiments and analysis. Q.Z. and C.D. performed and interpreted the structural modelling of variant EZH2 proteins. M.B. and J.A.M. performed and interpreted the extent of disease clustering and ΔΔG analyses of pathogenic EZH2 mutations. O.D., E.T. and A.P.B. co-wrote the manuscript with contributions from all other authors.

---

**Supplemental Figure S1.**
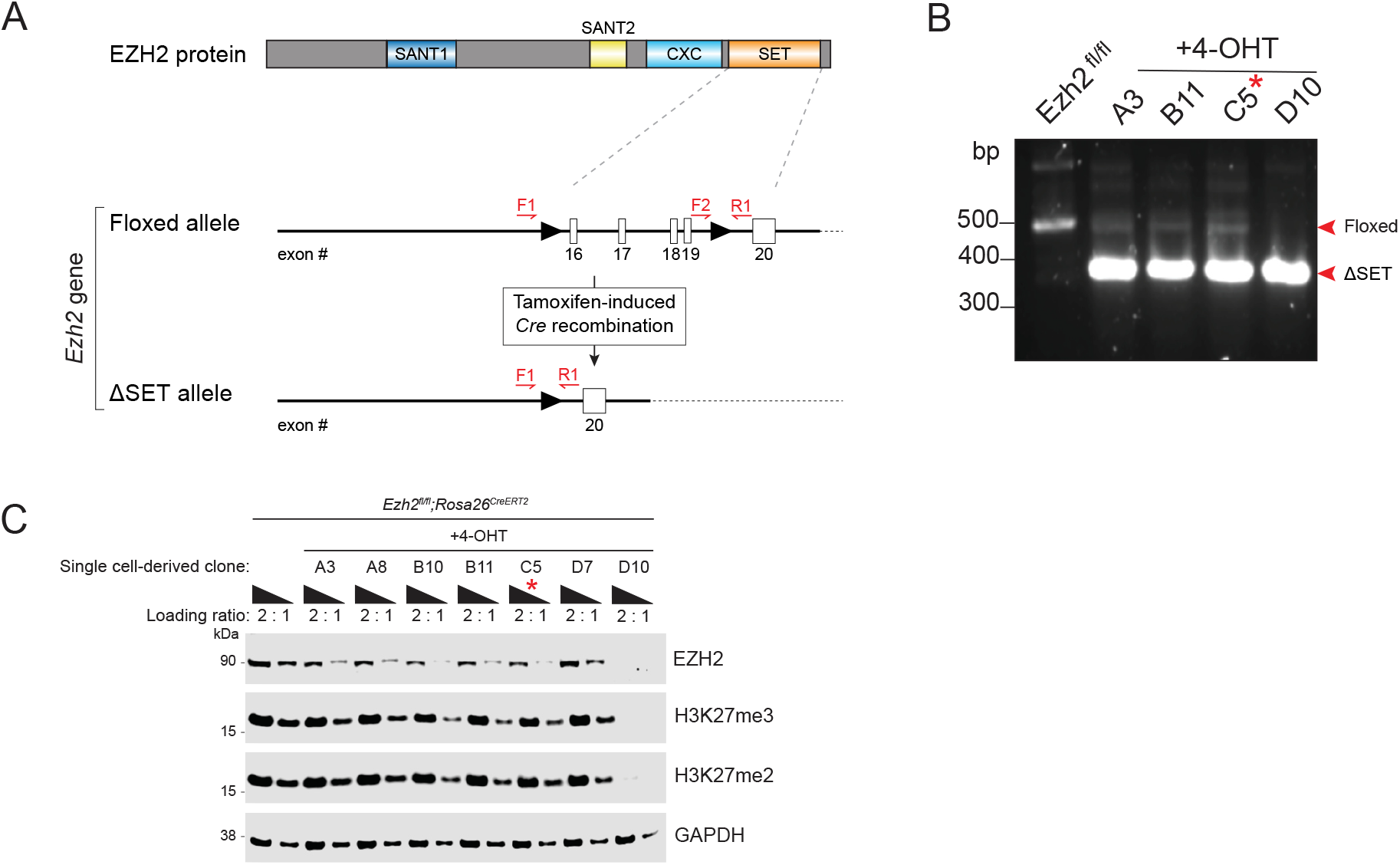
Generation of *Ezh2* heterozygous knockout mouse embryonic stem cells. **A**. (Top) Simplified schematic of the mouse EZH2 protein showing key domains. (Bottom) Schematic representation of the *Ezh2*-targeted locus before and after Cre-mediated recombination. SET domain-encoding exons are numbered and displayed as open boxes, while *loxP* sequences are depicted by arrows. Genotyping PCR primers are indicated in red. This gene targeting strategy was originally devised by the Tarakhovsky lab (Su et al. 2003). **B**. A representative genotyping PCR agarose gel using the primers indicated in (a), confirming heterozygous (A3, B11, C5) or homozygous (D10) knockout of the *Ezh2* SET domain-encoding exons. The asterisk indicates the clone (C5) that was taken forward as the parental *Ezh2*^*fl/Δ*^ cell line for all subsequent experiments. **C**. Immunoblots for EZH2, H3K27me3 and H3K27me2, probing whole cell lysates from the indicated cell lines. GAPDH is shown as a loading control. The asterisk indicates the clone (C5) that was taken forward as the parental *Ezh2*^*fl/Δ*^ cell line for all subsequent experiments. *Abbreviations: het, heterozygote; WT, wild-type; 4-OHT, 4-hydroxytamoxifen*.

**Supplemental Figure S2.**
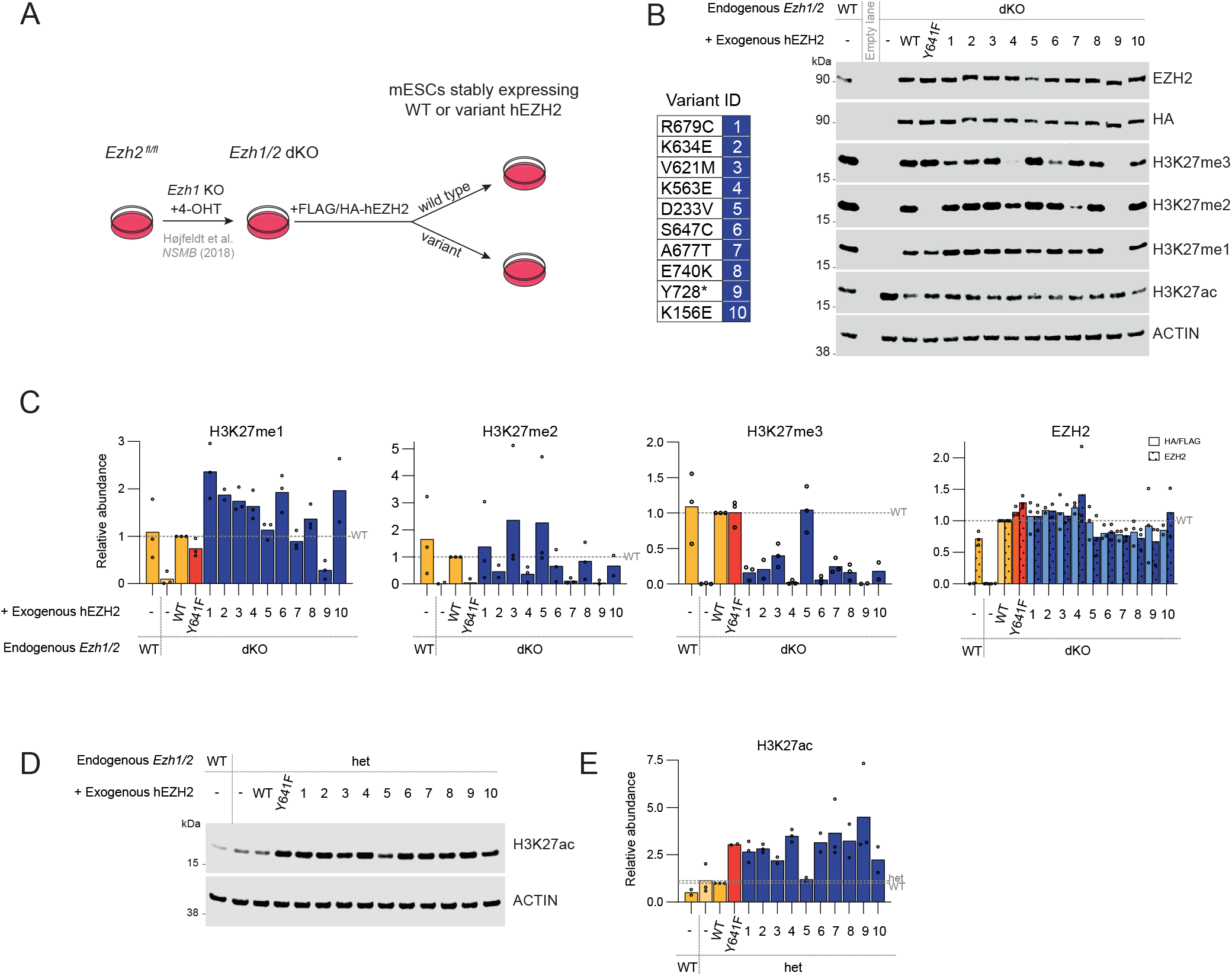
Weaver syndrome-associated EZH2 variants dominant negatively impair global PRC2 activitys. **A**. Schematic of the isogenic stem cell model system used to generate the data presented in panels b and c, involving expression of exogenous EZH2 constructs in *Ezh1/2* double knockout mouse embryonic stem cells. **B**. Representative immunoblots for HA, EZH2, and H3K27 modification states, probing whole cell lysates from the indicated cell lines. β-ACTIN is shown as a loading control. **C**. Relative quantification of immunoblot analyses of EZH2 and the HA/FLAG epitope tag, H3K27me1, H3K27me2 and H3K27me3 abundance in the indicated cell lines. Signal intensity values were normalized to a loading control and then expressed relative to the *Ezh1/2* dKO + EZH2-WT cell line. The dashed line represents the normalised *Ezh1/2* dKO + EZH2-WT value. Data values represent independent biological replicates whose median values are plotted as bars. **D**. Representative immunoblots for HA, EZH2, and H3K27 modification states, probing whole cell lysates from the indicated cell lines. β-ACTIN is shown as a loading control. **E**. Relative quantification of fluorescent immunoblot analyses of H3K27ac abundance in the indicated cell lines. Signal intensity values were normalized to a loading control and then expressed relative to the Ezh2^fl/Δ^ + hEZH2-WT cell line. The dashed lines represent the normalised Ezh2^fl/Δ^ (het) and Ezh2^fl/Δ^ + hEZH2-WT (WT) values. Data values represent n=2 or n=3 independent biological replicates whose mean values are plotted as bars. *Abbreviations: dKO, double knockout; het, heterozygote; WT, wild-type; 4-OHT, 4-hydroxytamoxifen*.

**Supplemental Figure S3.**
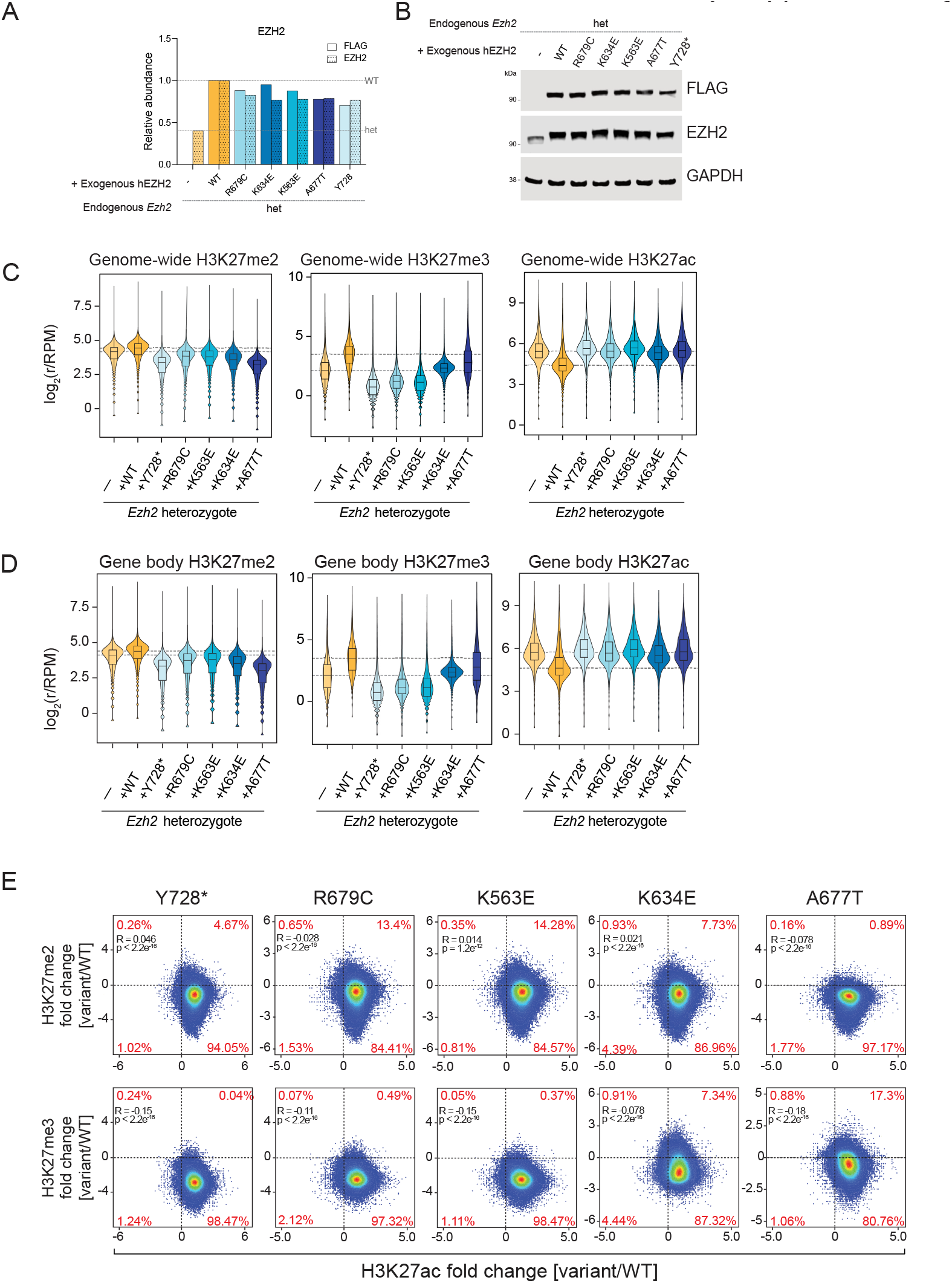
Weaver syndrome-associated EZH2 variants deplete intergenic H3K27 methylations and impair global chromatin compaction. **A**. Relative quantification of the ChIP-Rx matched immunoblot analysis of EZH2 and the FLAG epitope tag shown in panel b. Signal intensity values were normalized to a loading control and then expressed relative to the Ezh2^fl/Δ^ + hEZH2-WT cell line. The dashed lines represent the normalised Ezh2^fl/Δ^ (het) and Ezh2^fl/Δ^ + hEZH2-WT (WT) values. The oncogenic EZH2-Y641F variant was included as a positive gain-of-function control. **B**. Immunoblots for FLAG and EZH2, probing whole cell lysates harvested from the indicated cell lines in parallel with the ChIP-Rx experiment. GAPDH is shown as a loading control. The oncogenic EZH2-Y641F variant was included as a positive gain-of-function control. **C**. Violin plot representations of the total (genome wide) log_2_ ChIP-Rx normalized reads per million (r/RPM) of H3K27me2, H3K27me3 and H3K27ac enrichments in the indicated cell lines across 272,477 10kb bins. Control cell lines are coloured yellow while Weaver syndrome cell models are coloured blue. The dashed lines represent the *Ezh2*^*fl/Δ*^ + EZH2-WT values. **D**. Violin plot representations of the log_2_ ChIP-Rx normalized reads per million (r/RPM) of H3K27me2, H3K27me3 and H3K27ac enrichments in the indicated cell lines across 196,505 10kb bins representing all gene bodies. Control cell lines are coloured yellow while Weaver syndrome cell models are coloured blue. The dashed lines represent the *Ezh2*^*fl/Δ*^ + EZH2-WT values. **E**. Correlation plots demonstrating the relationship between changes in genome-wide ChIP-Rx enrichment of H3K27me2 (top row) or H3K27me3 (bottom row) and H3K27ac in cells expressing the indicated Weaver syndrome EZH2 variants compared to EZH2-WT. The percentage of 10kb bins in each quadrant is indicated. The p value and correlation coefficient were calculated using Pearson’s correlation. *Abbreviations: ChIP-Rx*, exogenous reference genome-normalised ChIP-seq; *het, heterozygote; r/RPM. ChIP-Rx normalized reads per million; WT, wild-type*.

**Supplemental Figure S4.**
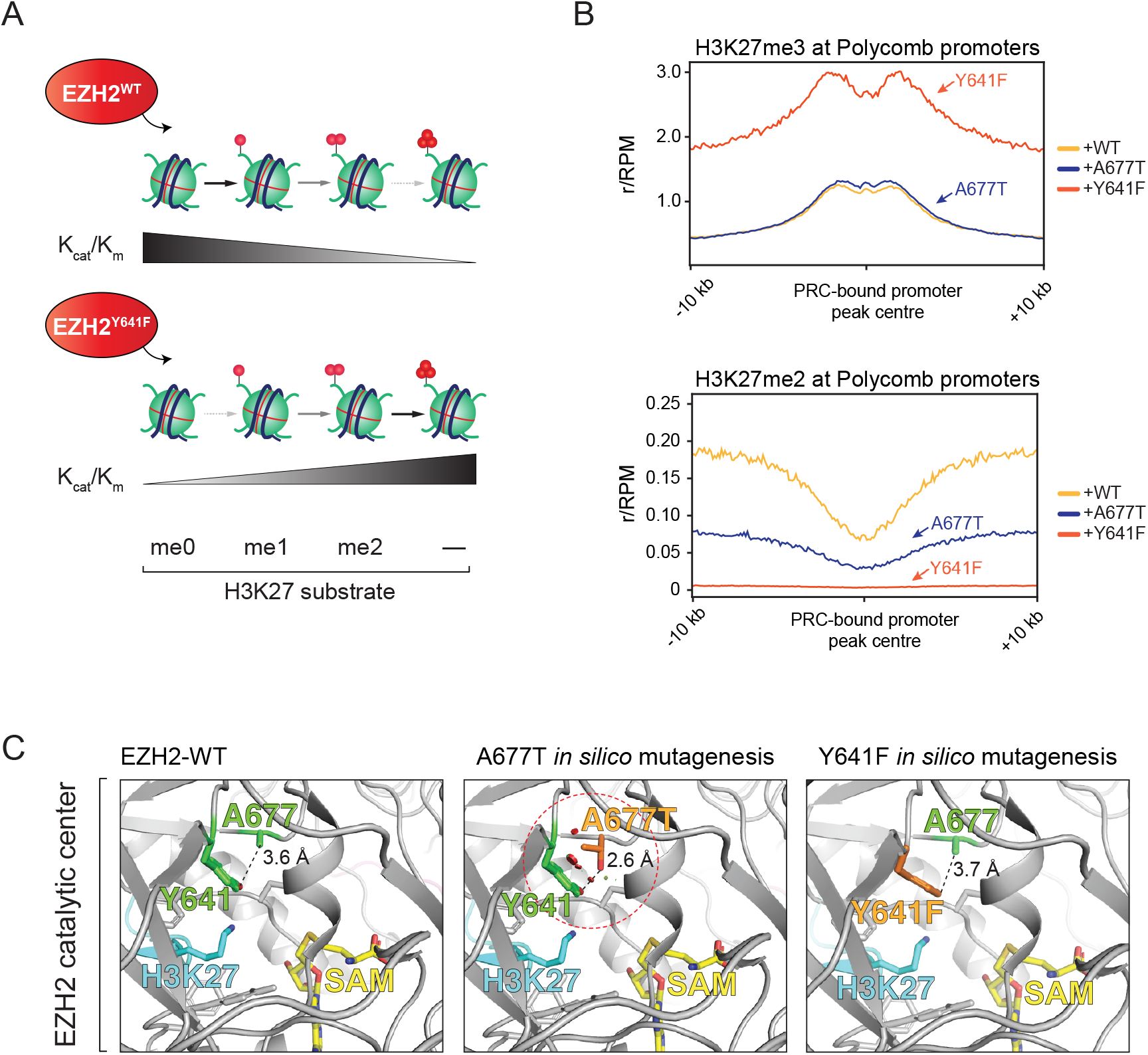
Weaver syndrome-associated EZH2 mutations differ in their effects on H3K27me3 at Polycomb target genes. **A**. Schematic illustrating PRC2-mediated H3K27me3 deposition at Polycomb target genes, where Kcat/Km indicates the catalytic efficiency of PRC2. **B**. Average ChIP-Rx signal profiles for H3K27me3 (top) and H3K27me2 (bottom) at PRC2-bound promoters (n = 3870) in Ezh2^fl/Δ^ mESCs expressing either EZH2-WT (yellow), EZH2-A677T (blue) or EZH2-Y641F (red). Data represents a biologically independent experiment (r2) from that presented in Figure 3C. **C**. Images of the *in-silico* mutagenesis of EZH2 to model the Weaver syndrome-associated hEZH2-A677T variant (middle) and the oncogenic hEZH2-Y641F (right) variant in the context of PRC2.2 with a substrate nucleosome (PDB 6WKR). The wild-type A677 and Y641 amino acids are coloured in green, with their corresponding mutated amino acids superimposed and coloured in orange. The substrate histone tail is in cyan, SAM is in yellow, and all other EZH2 residues are in grey. In the image of the A667T *in-silico* mutagenesis, steric (van der Waals) clashes between T677 and Y641 are represented by red discs (circled in red), while the dashed line indicates a short distance between the hydroxyls of both residues (2.6 Å). *Abbreviations: ChIP-Rx*, exogenous reference genome-normalised ChIP-seq; *het, heterozygote; r/RPM. ChIP-Rx normalized reads per million; WT, wild-type*.

**Supplemental Figure S5.**
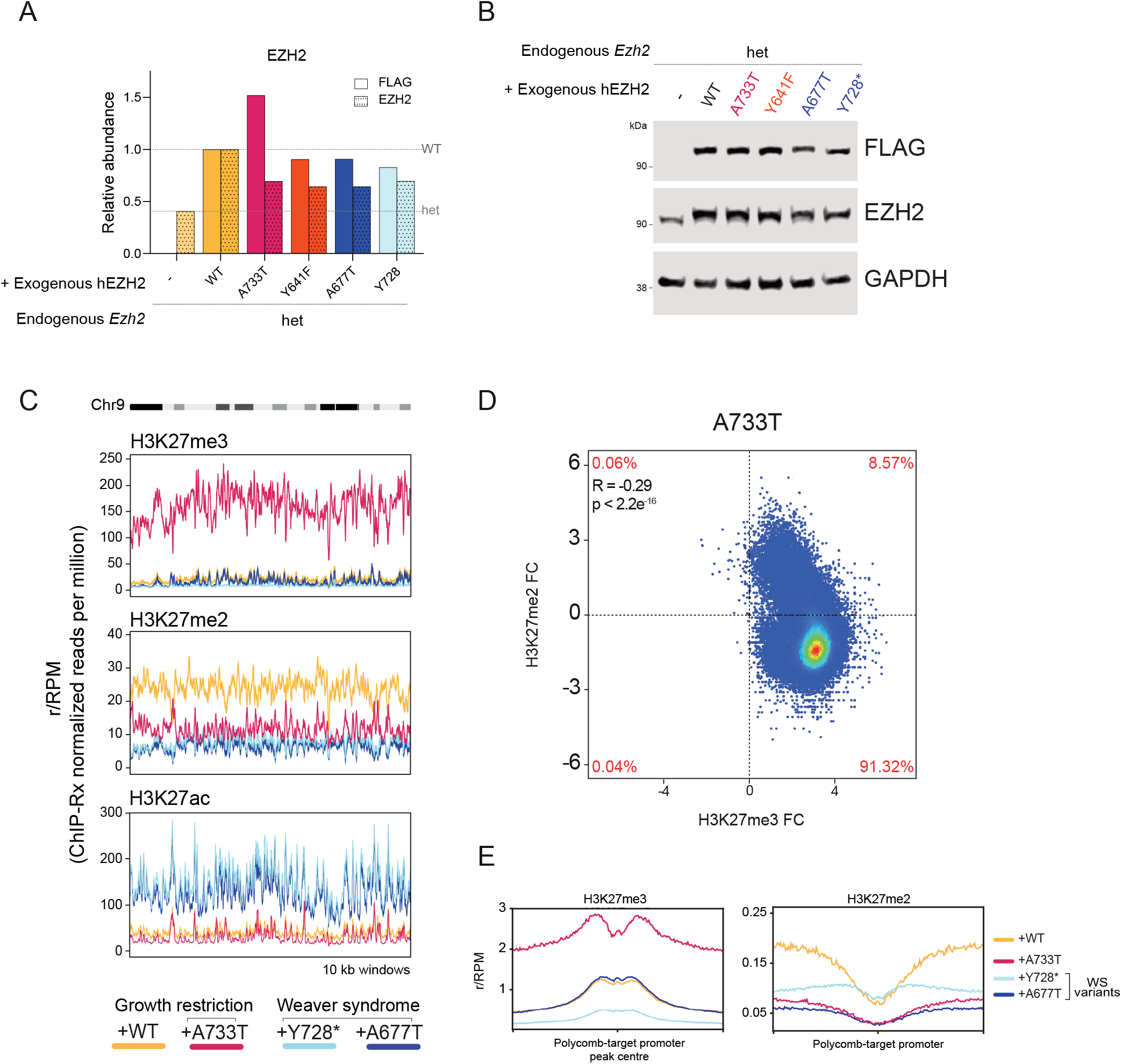
The growth restriction-associated EZH2-A733T variant induces opposing global H3K27 modification changes. **A**. Relative quantification of the ChIP-Rx matched immunoblot analysis of EZH2 and the FLAG epitope tag shown in panel b. Signal intensity values were normalized to a loading control and then expressed relative to the Ezh2fl/Δ + hEZH2-WT cell line. The dashed lines represent the normalised Ezh2fl/Δ (het) and Ezh2fl/Δ + hEZH2-WT (WT) values. **B**. Immunoblots for HA and EZH2, probing whole cell lysates harvested from the indicated cell lines in parallel with the ChIP-Rx experiment. GAPDH is shown as a loading control. **C**. Rolling average plots presenting the fold change in H3K27me3, H3K27me2 and H3K27ac ChIP-Rx signal in *Ezh2*^*fl/Δ*^ cells expressing growth restriction (pink) or Weaver syndrome (blue) EZH2 variants or wild-type EZH2 (yellow) across the whole of chromosome 9. **D**. Correlation plot demonstrating the relationship between changes in genome-wide ChIP-Rx enrichment of H3K27me2 and H3K27me3 in the growth restriction-associated EZH2 variant cell line, compared to wild-type EZH2. The percentage of 10kb bins in each quadrant is indicated. The p value and correlation coefficient were calculated using Pearson’s correlation. **E**. Average ChIP-Rx signal profiles for H3K27me3 and H3K27me2 at Polycomb target promoters (n = 3870) in the indicated cell lines. *Abbreviations: ChIP-Rx*, exogenous reference genome-normalised ChIP-seq; *het, heterozygote; r/RPM. ChIP-Rx normalized reads per million; WT, wild-type*.

